# Genotype imputation and error estimation in connected multiparental populations

**DOI:** 10.1101/2025.10.09.681496

**Authors:** Chaozhi Zheng, Eligio Bossolini, Nataša Formanová, Antje Rohde, Martin P. Boer, Fred A. van Eeuwijk

## Abstract

Multiparental populations have been produced for quantitative trait loci (QTL) mapping in many crops, where next-generation sequencing has become a cost-effective tool for genotyping. Previously, we have developed a hidden Markov framework denoted by MagicImpute_mma for genotype imputation in a multiparental population, which was implemented in Mathematica. However, its computational time increases quickly with the number of parents. In this work, we extend MagicImpute_mma into MagicImpute for increasing computational efficiency and robustness to various types of errors. Particularly, it has the following novel features: (1) allowing for multiple multiparental populations that may be connected by sharing parents, (2) allowing for many missing parents that are not available for sequencing, (3) accounting for allelic bias and overdispersion in next generation sequencing data, (4) inferring marker-specific error rates and filtering for markers with low error rates, and (5) being implemented in the high performance Julia language. Besides extensive simulation studies, we evaluate MagicImpute by three real datasets: the rice F2 population with sequence depth being low, the apple F1 population with parents being outbred, and the sorghum multi-parent advanced generation inter-cross (MAGIC) population with 10 male sterile lines (out of 29 parents) being missing. The results have shown that MagicImpute is accurate for genotype imputation in connected bi- or multiparental populations with various types of sequence errors and it opens up new opportunities for QTL mapping after imputing many missing parents.

## Introduction

Multiparental populations are experimental crosses usually produced by first outcrossing many inbred or outbred parents, then intercrossing systematically or randomly for some generations, and finally and optionally inbreeding by several generations of selfing or sibling mating. In comparison to traditional biparental populations, multiparental populations have high genetic diversity, and thus many such populations have been recently produced for QTL mapping in crops such as maize (Dell’acqua *et al*. 2015) and sorghum (Ongom and Ejeta 2018). For a given species, particularly in commercial-scale plant breeding, a large number of bi- or multi-parental populations are sometimes available and they are often connected by sharing parents. The focus of this work is methodology development for genotype imputation in connected bi- or multi-parental populations.

Genotype imputation methods have been originally developed for applications in human genetics, which can be generally categorized into population-based imputation using linkage disequilibrium information and pedigree-based imputation using linkage information. However, these methods are not optimized for experimental crosses in plant breeding. For example, the pedigree-based methods such as Merlin (Abecasis *et al*. 2002) and GIGI (Cheung *et al*. 2013) are not efficient for breeding pedigrees with selfing and ungenotyped members in intermediate generations, whereas the population-based methods such as Beagle5 (Browning and Browning 2016) and IMPUTE2 (Howie *et al*. 2009) are tailored to work with a reference haplotype panel and they cannot utilize pedigree information.

There are several existing methods for imputation in biparental populations, including FSFHap (Swarts *et al*. 2014), LB-Impute (Fragoso *et al*. 2016), and AlphaFamImpute (Whalen *et al*. 2020). In comparison, methods for multiparental populations are relative few, including mpimpute (Huang *et al*. 2014), R/qtl2 (Broman *et al*. 2019), and MagicImpute_mma(Zheng *et al*. 2018). However, mpimpute and R/qtl2 work only for specific mating designs and do not allow for missing data in parents. Although MagicImpute_mma has no such limitations, it is not computationally efficient in the case of many parents. Specifically, MagicImpute_mma performs founder imputation by analyzing all possible founder haplotypes along chromosomes, and thus the computational time increases quickly with the number of missing founder genotypes at each marker.

The primary aim of this work is to extend MagicImpute_mma (Zheng *et al*. 2018) for increasing computational efficiency in the case of many parents. Besides implementing MagicImpute in the high performance Julia language (Bezanson *et al*. 2017) instead of Mathematica (Wolfram Research 2016) for MagicImpute_mma (Zheng *et al*. 2018), we extend founder imputation algorithm for connected multiparental populations, particularly allowing for some or all founders’ genotypes being completely missing. Specifically, we partition founders into blocks and impute founder blocks alternatively.

The second aim of this work is to increase imputation robustness and heterozygous genotype calling accuracy for low coverage next generation sequencing data. Incorrect heterozygous calling can be due to low read depth and sequence errors. Allelic bias is a typical source of error, possibly because the polymerase chain reaction (PCR) amplification efficiency depends many factors such as GC content and the length of restriction fragments in sequence technologies (Schirmer *et al*. 2015; Salk *et al*. 2018; Qin *et al*. 2023). Genotype-Corrector (Miao *et al*. 2018) and GBScleanR (Furuta *et al*. 2023) have developed a binomial model for sequence data with the probability parameter accounting for allelic bias. We further develop a betabinomial model to account for both allelic bias and overdispersion.

This paper is organized as follows. We first describe MagicImpute algorithm for connected multiparental populations. Then we evaluate MagicImpute by extensive simulation studies, particularly testing the imputation of many ungenotyped founders and the estimation of various error rates. Furthermore, we evaluate MagicImpute by the real sorghum Multiparent Advanced Generation Inter-Cross (MAGIC) with 10 out of 29 founders being ungenotyped (Ongom and Ejeta 2018) and the two heterozygous populations: the rice F2 (Furuta *et al*. 2017) and the apple cross pollinated (CP) (Gardner *et al*. 2014) with low coverage sequences. In addition, we compare MagicImpute with GBScleanR (Furuta *et al*. 2023) using the rice F2. Finally, we discuss MagicImpute algorithm from the aspects of founder imputation and error estimation.

## Methods

### Overview of MagicImpute

The target population of MagicImpute is a population consisting of one or more subpopulations. Each subpopulation is derived from one or multiple founders via a few generations of mating crosses, and offspring in each subpopulation are exchangeable and thus they have the same prior genomic contributions from founders. A subdivided population is also termed connected populations if its subpopulations are connected by sharing founders. MagicImpute requires two input files: a variant call format (VCF) (Danecek *et al*. 2011) file for the genotypic data of founders and sampled offspring and a text file for the pedigree information.

Figure 1A shows the workflow of MagicImpute for a subdivided population, consisting of founder and offspring analyses. Marker binning is performed only if the average number of markers per bin is greater than the default 1.5, so that successive markers with distances less than the default 0.001 cM are binned. In the case of marker binning, the founder analysis is first performed for representatives and then for all markers. To increase the robustness of founder genotype estimation, the founder analysis is iterated several times until the run with the largest likelihood is close enough to the second best run so far.

**Figure 1.**
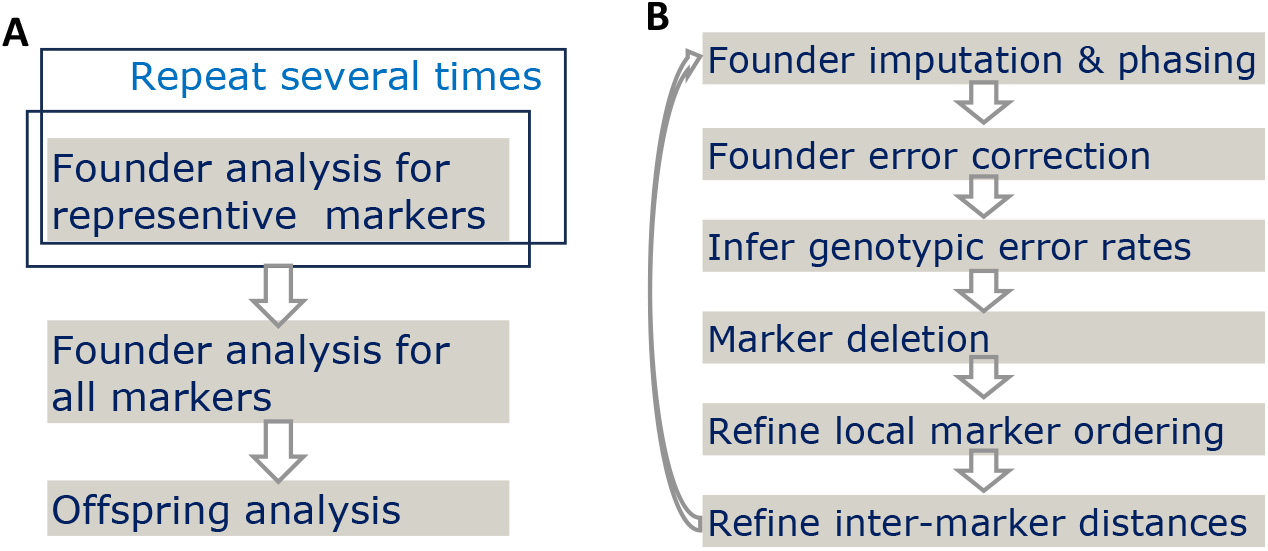
An overview of MagicImpute. (A) The workflow of MagicImpute. (B) The workflow of one-time founder analysis. The sub-steps are performed iteratively until all of them stop.

The MagicImpute algorithm builds on the hidden Markov model framework (Zheng *et al*. 2014, 2015, 2018), where the genotype model is a basic component. In the following, we first extend the data model of Zheng *et al*. (2018) to account for additional types of errors. Then we describe each sub-step of the founder analysis in Figure 1B. In comparison, MagicImpute_mma contains only the first sub-step. The offspring analysis has been described (Zheng *et al*. 2018) under the same assumption: offspring are independent conditional on the imputed phased founder genotypes.

### The genotype model

The genotype model calculates the probability of observed genotypic data given hidden states, which is also termed emission probability in the HMM. We assume markers are bi-allelic. For offspring *i* at marker *t*, we denote by *y*_*ti*_ the genotypic data, *ϵ*_*ti*_ the allelic error probability, and *z*_*ti*_ the hidden true offspring genotype. For founder *f* at marker *t*, we denote by 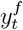 the genotypic data, 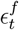 the allelic error probability, and 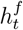 the hidden founder haplotype accounting for missing data and unknown phases but not allelic errors. In addition, let 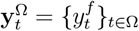 be the genotypic data for a subset Ω of founders at marker *t*, and 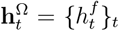 the hidden haplotypes, where Ω = *F* denotes the full set. We assume 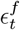 is a constant denoted by *ϵ*^*F*^ and set *ϵ*^*F*^ = 0.005 by default.

#### Called genotype

We first consider that *y*_*ti*_ is represented by a called genotype, corresponding to GT in the FORMAT field of a VCF file. The likelihood *P* (*y*_*ti*_|*z*_*ti*_, *ϵ*_*ti*_) is determined by an error model. We assume a random allelic error model: typing errors occur independent for each allele, and the resulting allele is the other if an error occurs. We obtain the marginal likelihood 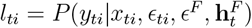 by summing out the hidden variable *z*_*ti*_

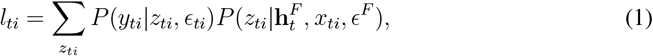

where *x*_*ti*_ is the hidden ancestral origin state, and the prior probability 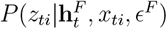 is given in Table S2 of Zheng *et al*. (2015).

To reduce the number of error parameters, the allelic error probability *ϵ*_*ti*_ is modeled in terms of marker-specific error probability *ϵ*_*t*_ and offspring-specific error probability *ξ*_*i*_,

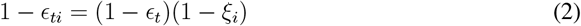

assuming that marker-specific errors occur independently of offspring-specific errors. The error probabilities *ϵ*_*t*_ and *ξ*_*i*_ will be estimated.

#### Allelic depth

We next consider that *y*_*ti*_ is represented by allelic depths in sequence data, corresponding to AD in the FORMAT field of a VCF file. We denote by *y*_*ti*_ = (*r*_0_, *r*_1_), where *r*_0_ and *r*_1_ are the numbers of reads for alleles 0 and 1, respectively. The read counts are modeled by the following binomial distributions

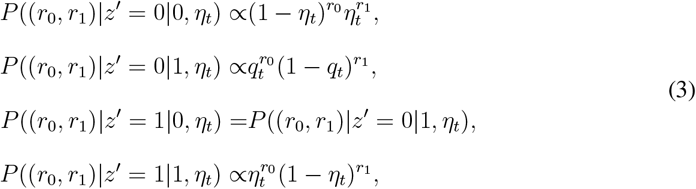

conditional on hidden true genotype *z*^*′*^ (Xie *et al*. 2010; Miao *et al*. 2018). Here phased genotype *z*^*′*^ is given in format of *a*_0_|*a*_1_, where *a*_0_ (or *a*_1_) denotes alleles 0 or 1. *η*_*t*_ denotes sequencing base error probability and *q*_*t*_ denotes allelic balance given the true *z*^*′*^ being heterozygous; *q*_*t*_ = 1*/*2 corresponds to no allelic bias.

To account for the variation of allelic balance among offspring, *q*_*t*_ is further modeled by a beta distribution

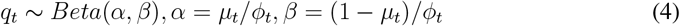

where shape parameters *α* and *β* are re-parameterized by mean *µ*_*t*_ and overdispersion *ϕ*_*t*_. Thus allelic depths *y*_*ti*_ is modeled by a beta-binomial distribution that approaches the binomial model with *q*_*t*_ = *µ*_*t*_ if *ϕ*_*t*_ goes to zero.

We introduce *ϵ*_*ti*_ to account for depth-independence errors such as misalignment, and the likelihood is given

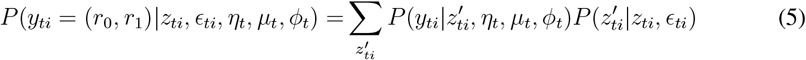

where *P* (*z′*_*ti*_|*z*_*ti*_, *ϵ*_*ti*_) is similar to *P* (*y*_*ti*_|*z*_*ti*_, *ϵ*_*ti*_) in equation (1) and *ϵ*_*ti*_ is given by equation (2). The final marginal likelihood is given by equation (1) with *P* (*y*_*ti*_|*z*_*ti*_, *ϵ*_*ti*_) replaced by *P* (*y*_*ti*_|*z*_*ti*_, *ϵ*_*ti*_, *η*_*t*_, *µ*_*t*_, *ϕ*_*t*_).

#### Prior distribution

At each marker locus, there are two error parameters *ξ*_*i*_ and *ϵ*_*t*_ for called genotypes, and five parameters *ξ*_*i*_, *ϵ*_*t*_, *η*_*t*_, *µ*_*t*_, and *ϕ*_*t*_ for allelic depths. We assign a weakly informative Beta prior distribution Beta(1.01, 1.01) for *µ*_*t*_ such that the distribution is symmetric at 0.5 (no allelic bias). We assign a Beta distribution Beta 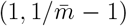 for each of the error parameters *ξ*_*i*_, *ϵ*_*t*_, and *η*_*t*_ and a prior exponential distribution with mean 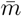 for *ϕ*_*t*_, where 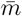 is the corresponding average over markers in the previous iteration during founder analysis.

### Founder imputation

To increase computational efficiency, we partition a set of founders into blocks such that each block must be a subset of the founders of a subpopulation and the block size must be no greater than a pre-specified upper bound *N*_*b*_. The founder partition is randomized in each iteration. We first sample a subpopulation and then sample a block of founders from the subpopulation without replacement, and repeat until there are no founders left. The sampling of a subpopulation is weighted by the total number of distinct offspring descended from the founders of the subpopulation.

The founder imputation starts from initializing founder haplotypes, where missing founder genotypes are randomly imputed and heterozygous genotypes for outbred founders are randomly phased. Specifically, let 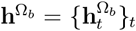 be the haplotypes for the *b*-th subset of founders at all loci. We impute 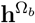 using the algorithm of Zheng *et al*. (2018), conditional on the current haplotypes in all the other blocks, and loop over each subset in each iteration. The founder imputation stops if the likelihood does not decrease for several successive iterations.

In each iteration, marker *t* is detected to be monomorphic in founders if there is only one allele in the current founder haplotypes 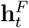. In initial five iterations, the observed founder genotypic data 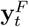 at each monomorphic marker *t* will be set to missing, aiming to reduce false detection because of errors in 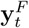. After five iterations, the monomorphic markers detected in each iteration are deleted.

### Founder error correction

In contrast to estimating the offspring error rates, we fix founder error rate *ϵ*^*F*^ to a small value (0.005 by default) and perform error correction. This is because offspring analysis is conditional on estimated founder haplotypes and remaining erroneous genotypes in founders can greatly decrease the estimation accuracy of offspring genotypes.

In each iteration, founder error correction is performed in several steps and it stops in the subsequent iterations if the number of corrections is zero. We first estimate the hidden ancestral origin states *x*_*ti*_ using the HMM Viterbi algorithm (Rabiner 1989; Zheng *et al*. 2015). Second, we obtain the best offspring genotypes by combining the estimated *x*_*ti*_ and the current founder haplotype **h**^*F*^. Third, we perform single site genotype calling if input data are allelic depths, and compare the results with the estimated best genotypes. Here single site genotype calling is based on equations (1) and (5), where markers are assumed to be independent. Lastly, at each marker for each founder we calculate the number of mismatches for each possible founder genotype, and perform correction if it reduces at least two mismatches.

Aiming to reduce false error corrections, the observed founder genotypic data corresponding to error corrections will be set to missing, such that these founder genotypes can only be imputed instead of being corrected in the following iterations in founder analysis (Figure 1B).

### Offspring error estimation

We first estimate the hidden ancestral origin states *x*_*ti*_ using the HMM Viterbi algorithm (Zheng *et al*. 2015; Rabiner 1989). Then we estimate each error parameter alternatively, conditional on all the other parameter values. For example, at each marker *t* we estimate *ϵ*_*t*_ by maximizing the conditional posterior distribution Π_*ti*_*p*(*ϵ*_*t*_), where *p*(*ϵ*_*t*_) is the prior distribution and *l*_*ti*_ is given by equations (1) and (5). Similarly, we estimate the other marker-specific parameters *η*_*t*_, *µ*_*t*_, and *ϕ*_*t*_ for allelic depths. Analogously, we estimate *ξ*_*i*_ by maximizing the posterior distribution Π_*ti*_*p*(*ξ*_*i*_).

We delete markers with too large error rates after five iterations. For each marker-specific error parameter, a marker is labeled if its error rate is larger than the corresponding hard threshold, or if its error rate is outlier and it is larger than the corresponding soft threshold. Similarly for the offspring-specific error parameter. The set of labeled markers is deleted, whereas the set of labeled offspring is temporarily excluded in the founder imputation and map refinement. The default hard thresholds for *ξ*_*i*_, *ϵ*_*t*_, *η*_*t*_, *µ*_*t*_, and *ϕ*_*t*_ are 0.25, 0.25, 0.05, 0.9, and 1.0, respectively; and the soft thresholds are 0.025, 0.025, 0.01, 0.67, and 0.01, respectively. An additional procedure for *µ*_*t*_ is performed to label markers with too small allelic balances. Outliers are detected based on the interquartile range using the Tukey’s fence 2 (Tukey 1977). According to our preliminary simulation studies, the commonly used fence 1.5 resulted in a few too many deletions, while the far outlier fence 3 resulted in too few deletions. The error estimation stops if the change of average error rates is very small.

In short, markers with an error rate being greater than its soft threshold are deleted only if they are also outliers, and thus the soft-threshold-based marker deletion can be disabled by setting Tukey’s fence to be infinitely large. In comparison, markers with an error rate being greater than its hard threshold are always deleted, and the hard-threshold-based marker deletion can be disabled by setting hard thresholds to be infinitely large.

### Marker deletion

Besides the marker deletion based on the estimated error rates, we perform marker deletion using the Vuong’s closeness test, a likelihood-ratio based test that can be used for comparing two non-nested models (Vuong 1989). Specifically, we first calculate the Vuong’s test statistic for each marker with the alternative hypothesis that the marker is removed. Then we remove markers using significance level 0.01. The level is somewhat arbitrary, and we did not observe a large effect of multiple testing in the simulation studies. The marker deletion stops if there is no marker deleted in the current iteration.

### Map refinement

The map refinement includes refining local marker ordering and inter-marker distances, which has been described in detailed by Zheng *et al*. (2019). It has been rearranged from the map construction package MagicMap (Zheng *et al*. 2019), so that the input marker map in MagicImpute can either be a physical map or be replaced by an optional map file. If the map file is provided by the output of MagicMap, it also contains a list of nearest markers for each marker. The neighborhoods can be used for neighbor-based order refining, which is much more effective than the default random order refining (Zheng *et al*. 2019). Thus, the order refinement is not performed by default if the optional map file is not provided.

### Evaluation by simulation

We evaluated MagicImpute by two groups of simulation studies. First, we simulated called genotypes in *n*-way RILs (n=2,4,8, and 16) by 6 generations of selfing, aiming to evaluate founder genotype estimation with or without repeating; each population was simulated with two sizes: 200 and 500. Second, we simulated sequence data in heterozygous mapping populations F2 and CP, aiming to evaluate the estimation of error parameters; each population was simulated with several sizes ranging from 5 to 1000. For both groups of simulations, we assumed five linkage groups with genetic length 100 cM for each. The number of markers per linkage group was set to 200 for the RIL populations, it ranged from 100 to 2000 for the heterozygous populations. The data simulation has been described in the previous studies (Zheng *et al*. 2015, 2018).

The simulation parameters for genotype missing were specified as follows. Independently for each marker, a true founder genotype was set to missing according to a Bernoulli distribution with probability parameter *p*, where *p* was sampled from beta distribution Beta(1, 19) with mean 0.05, and similarly for a true offspring genotype. For a RIL population with a given *n* founders and a given population size, we performed *n* + 1 independent simulations and set all the first *k* founders’ genotypes to missing in the (*k* + 1)-th simulation, where *k* = 0, …, *n*.

The simulation parameters for error rates were specified as follows. The genotypic error model in simulation was the same as the genotype model for imputation. For simplicity, offspring-specific error rate *ξ*_*i*_ was not simulated. At marker *t, ϵ*_*t*_ for offspring was sampled from Beta(1, 99) with mean 0.01, and similarly 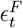 for founders was sampled from Beta(1, 99). For simulating sequence data, *η*_*t*_ was sampled from Beta(1, 499) with mean 0.002, *µ*_*t*_ was sampled from Beta(3, 3) with mean 0.5 (no allelic bias), and *ϕ*_*t*_ was sampled from an exponential distribution with mean 0.3. The Beta distributions for 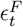, *ϵ*_*t*_, and *η*_*t*_ were set such that their means were twice as large as the default values of MagicImpute (see Table 1). In addition, the number *D*_*it*_ of reads (read depth) for offspring *i* at marker *t* was sampled from a Poisson distribution with mean *D*_*t*_, where *D*_*t*_ was sampled from an exponential distribution with mean being 5, 10, 20, and 40.

**Table 1.** List of nested error models for sequence data. The default values for parameters that are not estimated: *ϵ*_*t*_ = 0.005, *µ*_*t*_ = 0.5, *ϕ*_*t*_ = 0, *η*_*t*_ = 0.001, and *ξ*_*i*_ = 0.

| Model | Parameters to be estimated | Incremental parameter |
| --- | --- | --- |
| M0 | Null | Null |
| M1 | $\epsilon_t$ | $\epsilon_t$ marker-specific offspring error |
| M2 | $\epsilon_t, \mu_t$ | $\mu_t$ measure of allelic bias |
| M3 | $\epsilon_t, \mu_t, \phi_t$ | $\phi_t$ allelic overdispersion |
| M4 | $\epsilon_t, \mu_t, \phi_t, \eta_t$ | $\eta_t$ sequence base error |
| M5 | $\epsilon_t, \mu_t, \phi_t, \eta_t, \xi_i$ | $\xi_i$ individual-specific offspring error |

### Evaluation by real data

We evaluated the founder genotype imputation by the sorghum MAGIC (Ongom and Ejeta 2018) with 29 founders and 194 offspring, where 10 of 29 founders were unavailable for sequencing. The population was developed by crossing 19 founders with 10 random sterile plants out of a genetic male sterile source, followed by 10 generations of random mating and then 7 generations of selfing. Unexpectedly, around 4% genotypes were heterozygous, and they were set to missing in our study. The genotypic data were filtered for markers with minor allele frequency MAF ≥ 0.05, founder missing fraction *p*_*f*_ *<* 0.8, and offspring missing fraction *p*_*o*_ *<* 0.8, resulting in 44,767 markers. After filtering, the missing fraction in the 19 genotyped founders was 0.11. Since the breeding pedigree was unknown, MagicImpute used the prior average number 11 of recombination breakpoints per sampled offspring per Morgan as an input, according to the mating design.

In addition, we examined a series of nested error models (Table 1) by stepwise inclusion of the inferring error parameters *ϵ*_*t*_, *µ*_*t*_, *ϕ*_*t*_, *η*_*t*_, and *ξ*_*i*_. We evaluated the impact of the error models by sequence data in two heterozygous biparental populations: rice F2 (Furuta *et al*. 2017) and apple CP (Gardner *et al*. 2014) with the number of offspring being 813 and 87, respectively. The genotypic data for the rice F2 were filtered for markers with MAF ≥ 0.1 and *p*_*o*_ *<* 0.95, resulting in 8372 markers; and the genotypic data for the apple CP were filtered for markers with MAF ≥ 0.1 and *p*_*o*_ *<* 0.8, resulting in 48,713 markers.

MagicImpute was performed with and without inferring inter-marker distances for each error model. The input physical maps were transformed into genetic maps using the constant recombination rates 4.0 and 2.5 cM/Mbp for rice (Furuta *et al*. 2017) and apple (Gardner *et al*. 2014; Di Pierro *et al*. 2016), respectively. The imputation results were evaluated in two quantities: genotypic disconcordance was calculated by masking 10% input genotypic data with read depth ≥ 10 and using them as pseudo-true genotypes; the number of recombination breakpoints is estimated by haplotype reconstruction using the imputed genotypic data.

See supplemental File S1 for technical details on data preparation and analyses.

## Results

### Evaluate by simulated RILs

Figures 2 shows that founder imputation was quite accurate even when some or all founders’ genotypes were completely missing in the 16-way RILs of size 200. The founder genotypic disconcordance was close to zero when the number of missing founders was no larger than 5, and then it increased roughly with the number of missing founders (Figure 2A). The fluctuation was mainly because the one-time founder analysis algorithm was very greedy and usually finished within around ten iterations such that it sometimes converged to local maximums. Further examinations showed that the founder disconcordances often occurred in blocks, and these haplotype blocks were often switched with each other. Repeating founder imputation several times reduced the magnitude and fluctuation of disconcordances at the cost of computational time (Figure 2B-D).

**Figure 2.**
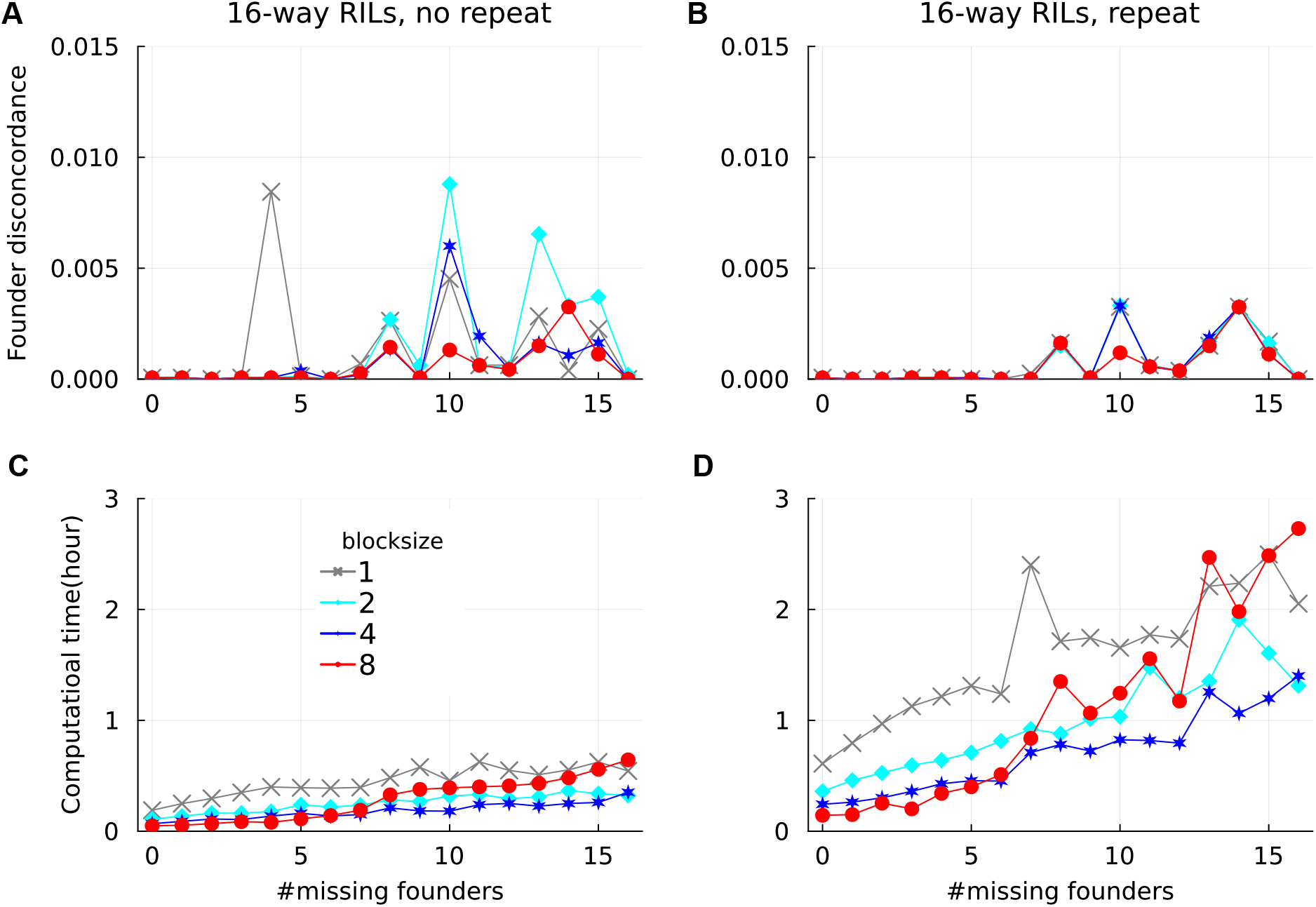
Founder imputation in the simulated 16-way RILs with population size 200. The x-axis denotes the number of founders with all genotypes being missing. The left and right panels show the results without and with repeating founder analyses, respectively.

The founder genotypic disconcordance decreased with population size. As shown in Figure S1 for the 16-way RILs, the founder disconcordances for population size 500 were generally smaller than those for population size 200, and they were mostly zero when repeating founder imputation. Whether founder imputation was repeated or not, the founder disconcordances for the *n*-way RILs with *n* ≤ 8 were mostly zero except for block size *N*_*b*_ = 1. Therefore, repeating founder imputation is not very necessary if the size of each subpopulation is large with respect to the number of founders for the subpopulation.

The founder genotypic disconcordance depended little on block size *N*_*b*_ (Figures 2 and S1), except that *N*_*b*_ = 1 often did not work well for bi-parental populations (results not shown). However, the computational time decreased with *N*_*b*_ up to 4, and for *N*_*b*_ = 8 it increased quickly when the number of missing founders was larger than 6 (Figures 2C-D and S1C-D). By default, MagicImpute set *N*_*b*_ = 2 for outbred founders and *N*_*b*_ = 4 for inbred founders.

### Evaluate by simulated F2 and CP

The estimation accuracy of the error rates, the correlation between estimated and true values, generally increased with read depth and population size (Figure 3). As shown in Figure S2, the estimation bias, the average difference between estimated and true values, decreased asymptotically into zero with population size and read depth. As expected, the estimations in small populations of size *<* 100 were not accurate and biased. The error estimations for the F2 were more accurate and less biased than that for the CP, probably because the latter retained more erroneous founder genotypes after imputation (Figure 4).

**Figure 3.**
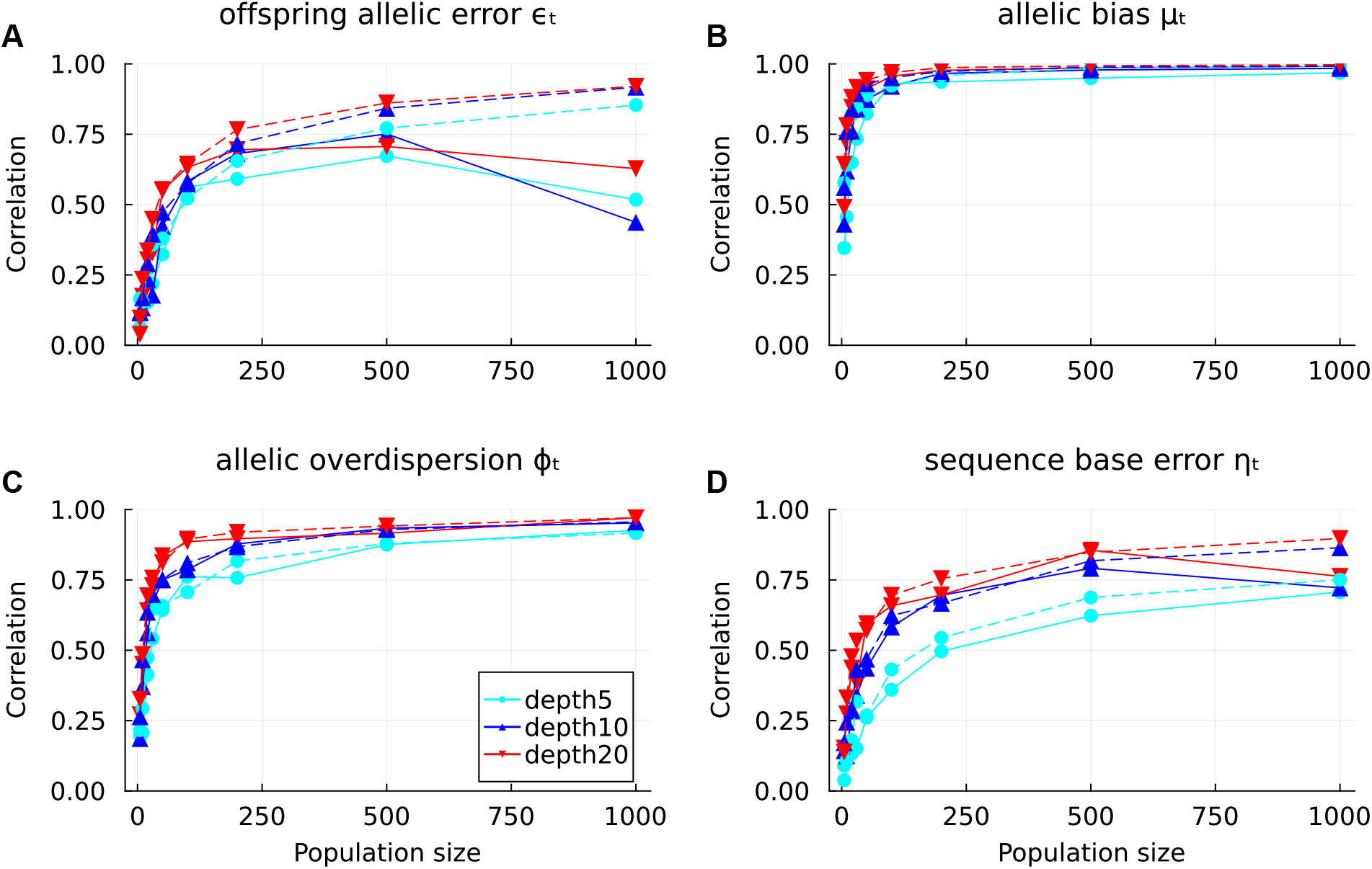
Estimation accuracy of error rates in the simulated F2 (dashed lines) and CP (solid lines). The y-axis indicates the correlation between estimated and true values among markers.

**Figure 4.**
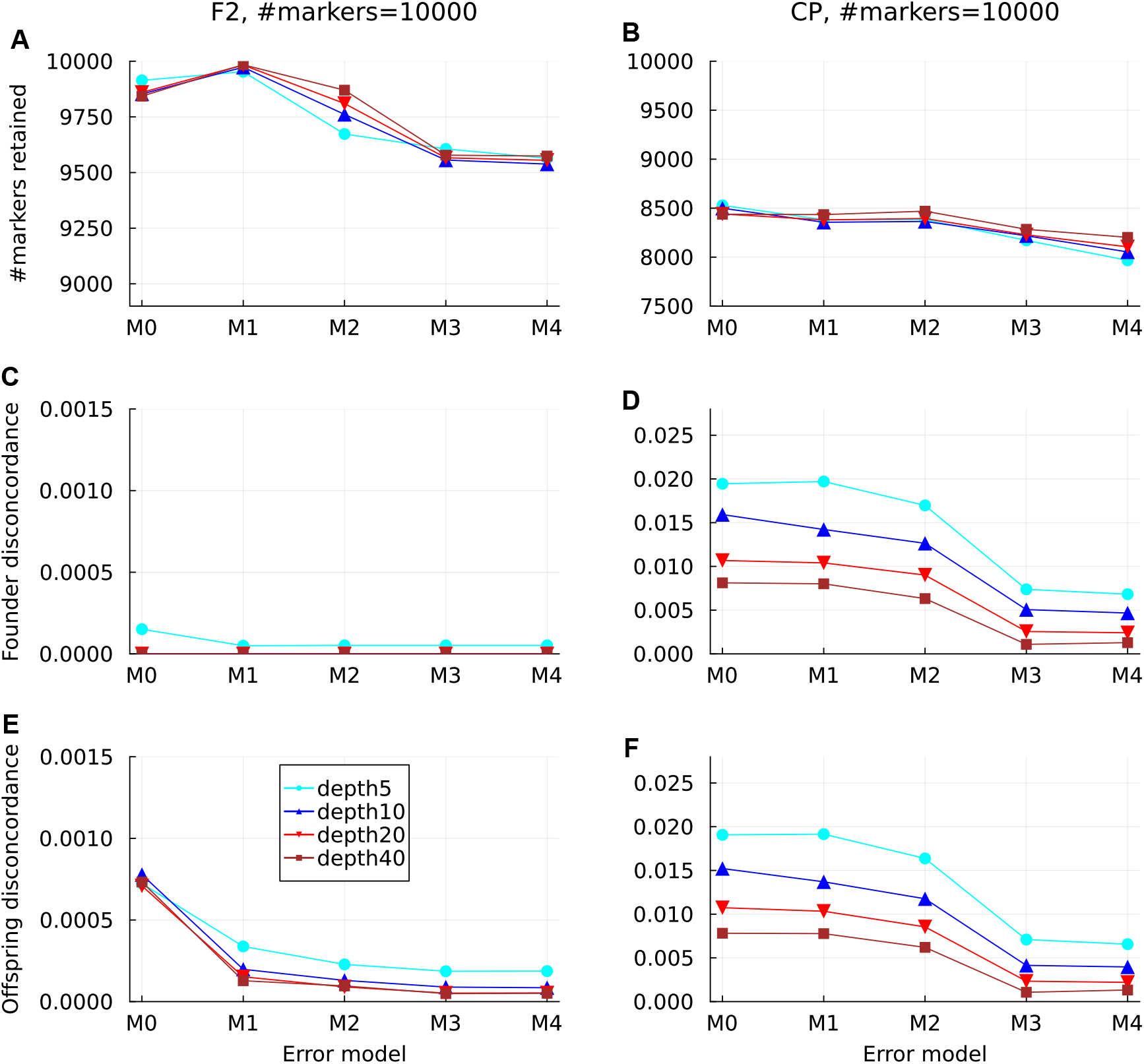
Genotype estimation in simulated F2 and CP. The results were obtained with the default founder error correction and the soft- and hard-threshold-based marker deletion. The error models are defined in Table 1. The disconcordance refers to the fraction of estimated genotypes being different from true values.

There were three actions affecting the impact of the nested error models (Table 1) on genotypic disconcordance: the founder error correction, the soft-threshold-based marker deletion, and the hard-threshold-based marker deletion. Figure 4 shows the results with all the three default actions, and Figures S3, S4, and S5 show the results obtained by forward stepwise disabling the actions. In the simulated F2, the hard-threshold-based deletion removed almost all founder genotypic errors, and the subsequent soft-threshold-based deletion and founder error correction were ineffective. In the simulated CP, the hard-threshold-based deletion removed roughly half founder genotypic errors (except null model M0); the subsequent soft-threshold-based deletion greatly reduced genotypic errors for models M3-M4, but they had essentially no effect for M0-M2; and the further founder error correction reduced genotypic errors for all the models, meanwhile it increased the number of retained markers. Briefly, a larger error model resulted in smaller genotypic disconcordances by deleting more markers with large estimated error rates.

On the other hand, the nested error models had negligible impact on the estimated number of recombination breakpoints, except that the null model resulted in more breakpoints; see Figure S6 with all the three actions. The estimated numbers of breakpoints for the non-null error models were almost independent of read depth (Figure S6A-B), but they asymptotically increased with the number of markers to the prior expected values (Figure S6C-D). Stepwise disabling the three actions resulted in no noteworthy differences with Figure S6.

### Evaluation by sorghum MAGIC

The imputation accuracies for the 19 genotyped founders ranged from 0.65 to 0.97 with mean 0.85, whereas the imputation accuracy for offspring was 0.993. After imputation, the average missing fractions was 0.25 for the 10 ungenotyped founders and 0.068 for the other 19 founders (Figure 5A), whereas it was 0.018 for the offspring. The missing genotypes in the 10 ungenotyped founders did not occur at random but in segments, probability because these segments were not inherited by any offspring.

**Figure 5.**
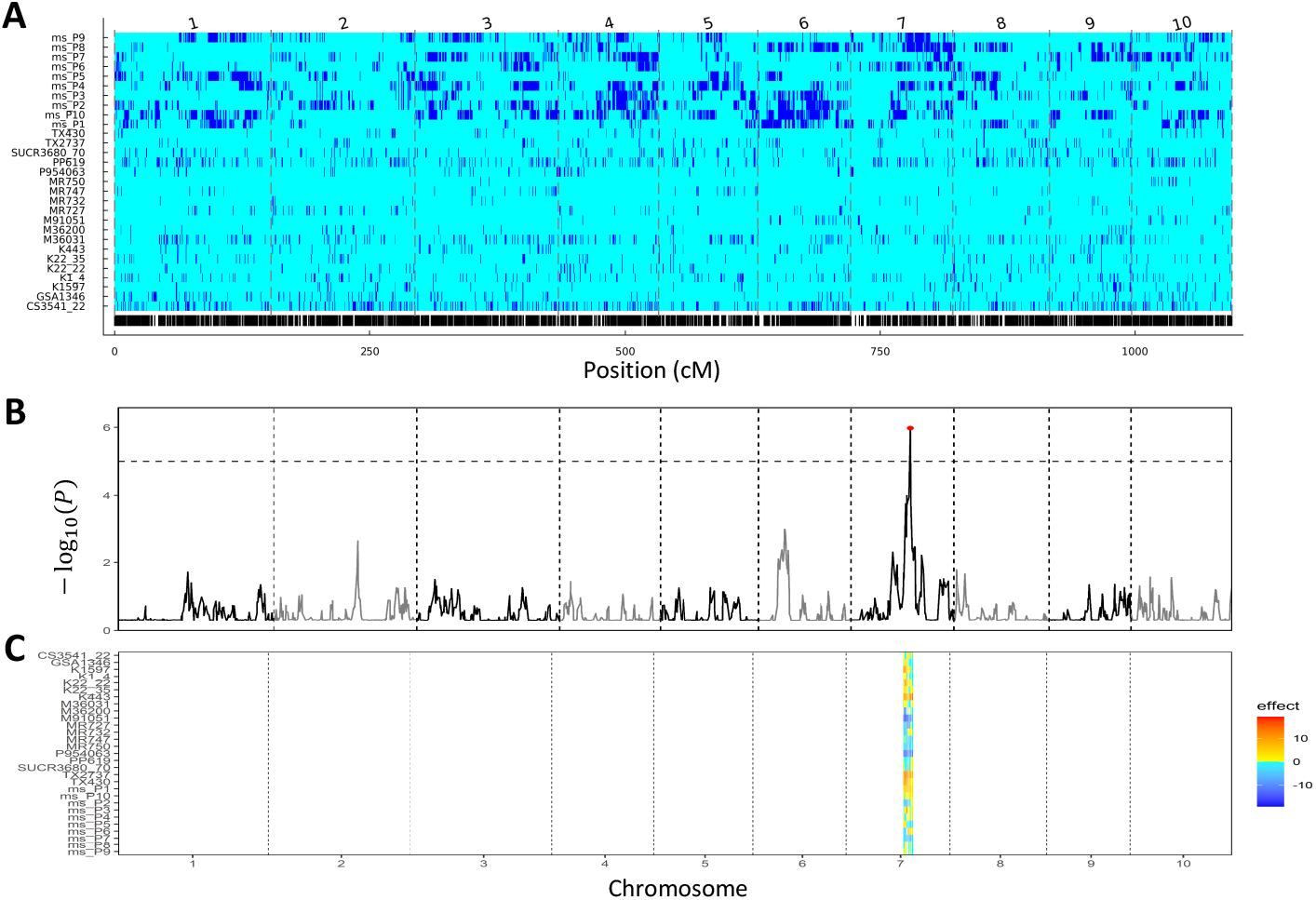
Founder imputation in the real sorghum MAGIC. The dashed vertical lines indicate the chromosome boundaries. (A) The missing pattern of founder genotypes after imputation. Blue and aqua denote missing and non-missing, respectively. The founders from ms_P1 to ms_P10 were completely missing before imputation. The bottom vertical lines denote marker positions. (B) QTL profile of plant height based on the posterior ancestral probabilities calculated from the imputed genotypic data. The horizontal dashed line denote the threshold. (C) The effect of founder origins at detected QTLs.

The large fraction of non-imputed founder genotypes had little effects on the estimated number of recombination breakpoints per offspring. The estimated genetic length was 1094 cM (Figure S7A), slightly smaller than the previous estimations ranging from 1192 to 1369 cM (Jin *et al*. 2021; De Souza *et al*. 2021; Somegowda *et al*. 2022; Guden *et al*. 2023). In return, the estimated average breakpoint density was 13.3 per Morgan per offspring, slightly larger than the prior expected value 11 (Figure S7B).

We further evaluated imputation by QTL mapping for plant height. The QTL profile in Figure 5B was obtained using the statgenMPP package (Li *et al*. 2022) with inputs being the posterior ancestral probabilities for all offspring and the average plant heights between years 2013 and 2014. In comparison to the previous GWAS (Ongom and Ejeta 2018), the major QTL in chromosome 7 was recovered, whereas the two other minor QTLs in chromosomes 6 and 9 were absent in current study (Figure 5B).

There are several possible reasons for the differences between current QTL mapping and the previous GWAS. The latter was based on the plant heights in the two years rather than the average values, whereas the statgenMPP worked only for a single environment. More likely, the population size 194 is not sufficient for accurately imputing the 29 founders, as shown in Figure 5A. On the other hand, founder imputation allows for QTL mapping that in turn allows for estimating the effects of founder origins (Figure 5C).

### Evaluation by rice F2 and apple CP

Figure 6 shows the effects of the nested error models (Table 1) in the real rice F2 and the apple CP. The genotypic disconcordances generally decreased with error model size except for M0 (Figure 6A-D), although the decreases were marginal for the rice F2. In contrast with the simulation studies, the number of recombination breakpoints decreased with error model size (Figure 6E-F). As shown in Figures S8 and S9, the excessive breakpoints mostly resulted from tiny segments, which might explain that the error models had only small effects on imputation accuracy.

**Figure 6.**
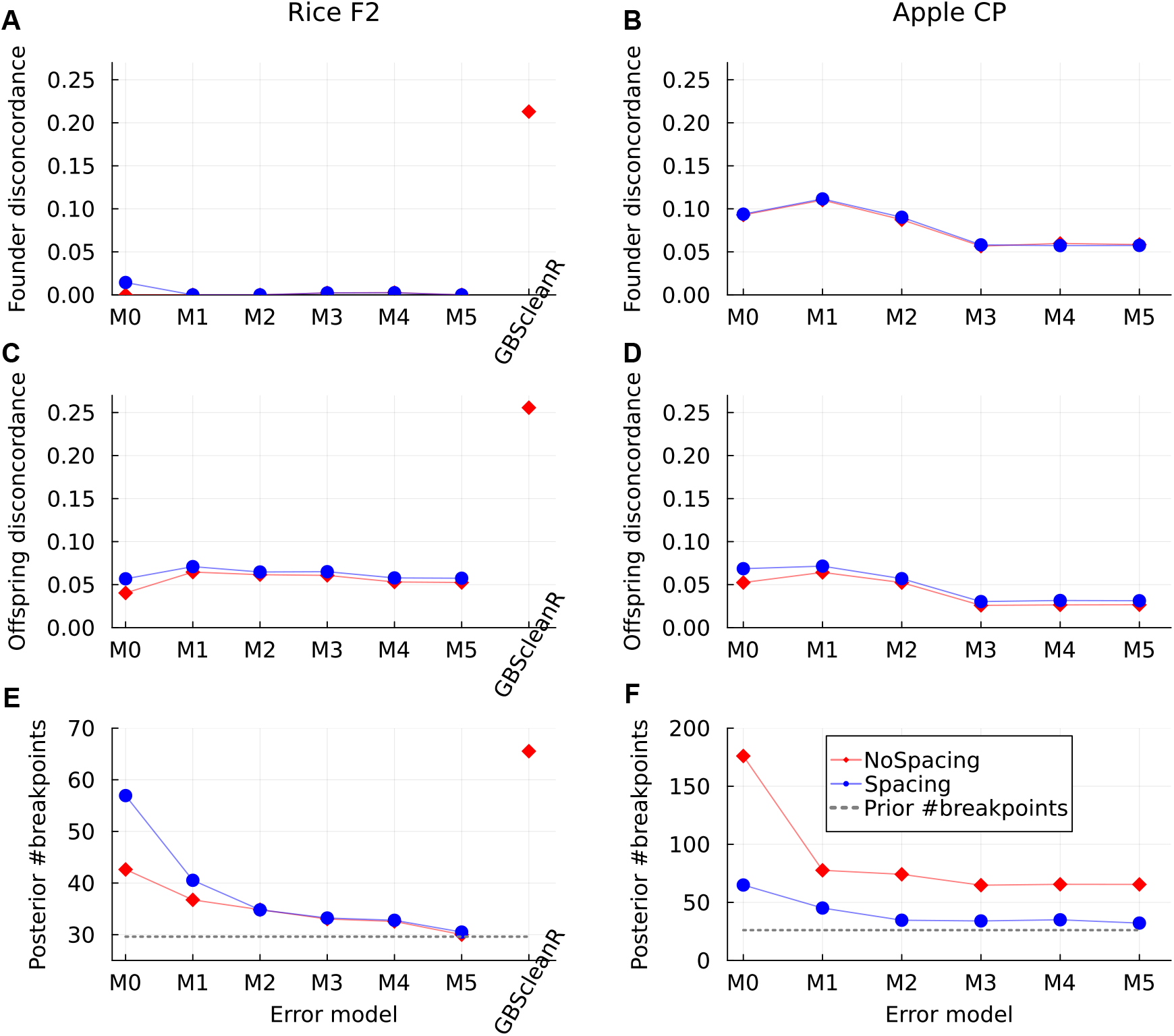
Impact of error models in the real rice F2 and apple CP. The error models are defined in Table 1. The blue and red lines denote the results with and without inferring inter-marker distances, respectively. The dotted lines in the bottom panels denote the prior number of recombination breakpoints per offspring.

To account for recombination rate varying along chromosomes, inter-marker distances were inferred during genotype imputation. As shown in Figure 6, it resulted in longer genomic lengths and more recombination breakpoints, but had little effects on genotypic disconcordance. The genomic length and the number of breakpoints decreased generally with error model size, and they leveled off at the expected values in the rice F2 but not in the apple CP (Figure 6E-F). This is probably because the input physical map for the apple CP has low quality (Di Pierro *et al*. 2016) and the errors due to marker misordering played a dominant role (Figures 6E-F)

In the rice F2 and the apple CP, there were typically around two markers in a chromosome stretch of length 75 bp. The genotypic disconcordances and the number of breakpoints became larger when nearby markers were not binned during imputation (Figure S11), indicating that nearby co-segregating markers had large influence.

As shown in Figures 6 & S11, MagicImpute obtained almost zero founder genotypic disconcordances by deleting around 45% erroneous markers during imputation, whereas GBScleanR retained all markers and resulted in much larger genotypic disconcordances (>0.2) and much more recombination breakpoints. In comparison, Furuta *et al*. (2023) obtained offspring disconcordance around 0.12 for the rice F2 using GBScleanR, probably because of stringent criteria of selecting markers for genotype masking (missing fraction < 0.2 and minor allele frequency > 0.4).

## Discussions

We have developed MagicImpute for genotype imputation in connected multiparental populations, especially increasing computational efficiency when there are a large number of founders. By partitioning founders into blocks, MagicImpute increases computation efficiency in two aspects. First, in each iteration the number of calculations for different founder haplotypes is much smaller than that of MagicImpute_mma, even though a few more iterations are required for MagicImpute. Second, conditional on the current founder haplotypes in each iteration, MagicImpute performs the standard HMM algorithms (Zheng *et al*. 2015; Rabiner 1989) for each subpopulation, in comparison that MagicImpute_mma analyzes the whole population jointly. The computational efficiency for MagicImpute is mainly determined by the block size *N*_*b*_ and the numbers of founders for each subpopulation, rather than the total number of founders for the whole population for MagicImpute_mma.

MagicImpute is capable of imputing founders even when some or all founders’ genotypes are completely missing, as shown in the simulation studies. Few imputation methods have such capacity. AlphaFamImpute (Whalen *et al*. 2020) works for full-sib families with two outbred ungenotyped parents, and MagicImpute_mma works only for a limited number of ungenotyped founders. Assuming a genetically diverse population is derived a given number of generations ago, STITCH (Davies *et al*. 2016) works for low-coverage sequences without requiring a reference panel. However, it needs a high-quality reference assembly. It is promising to further extend MagicImpute for genetically diverse populations by imputing virtual founders.

The evaluation by the real Sorghum MAGIC has also shown the feasibility of QTL mapping after imputing ungenotyped founders, although a larger population size might be necessary for accurately imputing so many missing founders and possibly demonstrating the outperformance of QTL mapping. Compared to GWAS, QTL mapping can also provide extra information such as the identification of potentially causal QTL haplotypes. However, a sophisticated QTL modeling for traits in multi-environments in multiparental populations is demanded.

MagicImpute has implemented various error models for sequence data, where each error parameter can be specified whether it is to be inferred or to be fixed to the default value. Our evaluations by simulation studies and real data have shown a larger error model can increase imputation accuracy by deleting more erroneous markers. Miao *et al*. (2018) have shown that Genotype-Corrector has larger correct rates of heterozygous genotype call in F2 populations than those of Beagle and LinkImpute (Money *et al*. 2015). In comparison, GBScleanR (Furuta *et al*. 2023) retained all markers and resulted in low imputation accuracy.

In contrast to the consistent effects on imputation accuracy, the effects of error models on the estimated number of recombination breakpoints are different between the simulation studies and the evaluations by real data. The HMM framework of MagicImpute assumes independence of genotypic data, conditional on hidden ancestral states; and the differential effects are very likely because the assumption of conditional independence is valid in the simulation studies, but it is violated for the real data. In the rice F2 and the apple CP, many reads span multiple markers within short sequence stretches. Apparently, the non-independence of reads can be partly accounted for by increasing the number of error parameters, and partly accounted for by marker binning in MagicImpute. A further extension to replace marker bins with multi-allelic markers instead of representative markers is under development.

MagicImpute is also suitable for constructing a consensus genetic map in connected mapping populations, assuming the input physical map has high quality. The inflation of genomic length can be alleviated by marker deletion and error estimation; particularly, the individual specific error rate *ξ*_*i*_ can account for different genotypic platforms and possible sample contamination. If the input physical map has low quality such as the apple CP, it is recommended to first construct a genetic map by the MagicMap (Zheng *et al*. 2019) and then refine it by MagicImpute.

In conclusion, MagicImpute is accurate for genotype imputation in connected multi-parental populations with a large number of subpopulations, and it is capable of imputing many missing founders provided there are a sufficient number of genotyped offspring.

## Data availability

MagicImpute is implemented as a package in the RABBIT software written in the Julia language (Bezanson *et al*. 2017). RABBIT is available from the web site: https://github.com/biometris/RABBIT. There are also command line interfaces for running RABBIT in a Windows or Linux shell. The data simulations were performed using MagicSimulate in the RABBIT, and the number of recombination breakpoints and the posterior ancestral probabilities were calculated from imputed genotypic data using MagicReconstruct in the RABBIT. The real data for the rice F2, the apple CP, and the sorghum MAGIC were downloaded based on the supplemental information of Furuta *et al*. (2017), Gardner *et al*. (2014), and Ongom and Ejeta (2018).

## Acknowledgments

This work was funded by BASF Belgium Coordination Center CommV. Author contributions: A.R. and F.A.E. contributed to conceptualization. C.Z. developed the algorithm and wrote the first draft of the manuscript. E.B. and N.F. have tested the software and provided valuable feedback. All authors provided critical comments and reviewed and edited the manuscript. All authors approved the final manuscript.

## File S1

### Details on data analysis

We performed data analyses using RABBIT v1.3.4; See the online manual https://github.com/Biometris/RABBIT for all RABBIT functions. For comparisons, we used GBScleanR v2.3.2 for analyzing the rice F2.

### Rice F2

The genotypic data file `“gbs_nbolf2.vcf.gz”` was downloaded online https://github.com/tomoyukif/GBSR_SupFiles. We analyzed the data by the following steps.

We first filtered the data by function `vcffilter` in the RABBIT

~~~
vcffilter(genofile;
 delsamples = [“03_F1”],
 maxmiss = 0.99,
 minmaf = 0.05,
 deldupe = true,
)
~~~

where option `delsamples` deletes the sample `“03_F1”`, `maxmiss` filters for markers with genotype missing fraction *<*= 0.99, `minmaf` filters for markers with minor allele frequency *>*= 0.05, and `deldupe` deletes successive markers having exactly the same genotypic data. After filtering, the number of markers was reduced from 20,224 to 11,398, and the filtered data was saved in `genofile2`.

We then filtered the new `genofile2` by function `magicfilter` in the RABBIT

~~~
magicfilter(genofile2,designcode;
 isfounderinbred = true,
 minmaf = 0.1,
 missfilter = (f,o) -> o < 0.95
)
~~~

where `designcode` is given by `“01_NB/02_OL=>1”`, denoting the two parents `“01_NB”` and `“02_OL”` were first crossed and the resulted F1 was then self-fertilized for one generation. Here `isfounderinbred` specifies whether parents are inbred, `minmaf` filters for markers with minor allele frequency *>*= 0.1, and `missfilter` filters for markers with genotype missing fraction in offspring *<* 0.95 regardless of missing in parents. The number of markers was further reduced to 8372, and a new `genofile3` and a pedigree file `pedfile` were output.

To calculate imputation accuracy, we masked the filtered data by function `magicmask` in the RABBIT

~~~
magicmask(genofile3,pedfile;
 isfounderinbred = true,
 minread=10
)
~~~

where `minread` denotes masked read depths must be no less than 10. By default, the read depths at each markers for each individual (founder of offspring) were masked with probability 0.1 if they met the `minread` requirement. The read depths were called into genotypes with threshold 0.9 and used as ground truths for calculating genotypic disconcordances. In the genotype calling, the default sequence error rates were used: no allelic bias, no overdispersion, and sequence base error is set to 0.001. A new `genofile4` was output.

Next, genotype imputation was performed by function `magicimpute` in the RABBIT

~~~
magicimpute(genofile4,pedfile;
 isfounderinbred = true,
 isphysmap = true, recomrate = 4,
 likeparam,
 threshlikeparam,
 softlikeparam,
 tukeyfence = 2,
 iscorrectfounder = true,
 isspacemarker = true,
 skeletonsize = 100,
)
~~~

where `isphysmap` and `recomrate` denotes transforming the input physical map into a genetic map with recombination rate 4 cM/Mbp, `iscorrectfounder` denotes whether to perform founder error correction, `isspacemarker` denotes whether to infer inter-marker distances, `tukeyfence` denotes Tukey’s fence 2, and `skeletonsize` denotes 100 skeleton markers being used for re-scaling estimated inter-marker distances. A new `genofile5` was output.

We set `likeparam = LikeParam(peroffspringerror=0.005, offspringerror = 0.005, allelicbias = 0.5, allelicoverdispersion = 0, baseerror = 0.001)` for null model *M* 0. The keywords `peroffspringerror`, `offspringerror`, `allelicbias`, `allelicoverdispersion`, and `baseerror` correspond to the symbols *ξ*_*i*_, *ϵ*_*t*_, *µ*_*t*_, *ϕ*_*t*_, and *η*_*t*_ in the body text, respectively. Setting the value of an error parameter to `nothing` will infer the error parameter. For example for error model *M* 3, `likeparam = LikeParam(peroffspringerror=0.005, offspringerror=nothing, llelicbias = nothing, allelicoverdispersion = nothing, baseerror = 0.001)`.

The option `threshlikeparam` specifices hard thresholds. By default, `threshlikeparam = ThreshLikeParam(peroffspringerror = 0.25, offspringerror = 0.25, allelicbias = 0.9, allelicoverdispersion = 1.0, baseerror = 0.05)`. The `softthreshlikeparam` specifices soft thresholds. By default, `softthreshlikeparam = SoftThreshLikeParam(peroffspringerror = 0.025, offspringerror = 0.025, allelicbias = 0.67, allelicoverdispersion = 0.01, baseerror = 0.01)`.

Finally, we run function `magicreconstruct` in the RABBIT

~~~
magicreconstruct(genofile5,pedfile;
  isfounderinbred = true,
  hmmalg = “viterbi”,
)
~~~

where `hmmalg` denotes using the Viterbi algorithm for haplotype reconstruction. The number of recombination breakpoints was calculated by the number of change-points in the inferred Viterbi path of ancestral origins for each offspring in each linkage group.

The `genofile4` output by `magicmask` was loaded by `GBScleanR` to produce object `gdata`. It was first filtered by

~~~
gdata <- setCallFilter(gdata,
  ref_qtile = c(0, read_quantile),
  alt_qtile = c(0, read_quantile))
~~~

where `read_quantile = 0.9` is used to discard read depths that are too high to be reliable. Genotype imputation was then performed by function `estGeno` in GBScleanR

~~~
gdate <- estGeno(object = gdata,
  recomb_rate = 0.04, iter = 4,
  error_rate = 0.0025,
  het_parent = FALSE,
  call_threshold = 0.9,
  optim = TRUE)
~~~

where `recomb_rate` is in unit of Morgan/Mbp so that the same value of 4 cM/Mbp is specified, `iter = 4` specifies the number of iterative parameter updates (2 by default), and the rest options are set by default.

### Apple CP

The apple CP data was analyzed using the same steps for the rice F2 except the following different option values.

Set `isfounderinbred = false` in functions `magicfilter`, `magicmask`, `magicimpute`, and `magicreconstruct`. Set `designcode` to `“Golden/Scarlet=>0”` for F1 cross without selfing. Set `delsamples = nothing` (no sample deletion) and `maxmiss = 0.9` in function `vcffilter`. Set `missfilter = (f,o) -> o <= 0.8` in `magicfilter` for filtering for markers with genotype missing fraction in offspring ≤ 0.8, since the apple CP has much smaller population size. Set `recomrate = 2.5` and `isrepeatimpute = true` in `magicimpute`; the default no repeating of founder analysis was used in the rice F2.

### Sorghum MAGIC

A genotypic data file in the VCF format was prepared based on the online supplemental data. The function `vcffilter` was not performed and `magicfilter` was performed with options `minmaf = 0.05` and `missfilter = (f,o) -> f < 0.8 && o < 0.8`. Since the breeding pedigree is not known, the population design was set as follow

~~~
sorghum_design=“nfounder=29||ibd=1.0||mapexpansion=11”
~~~

where the 29 founders must come before offspring in the VCF file, `ibd=1.0` denotes all offspring are assumed to be completely inbred, and `mapexpansion=11` denotes the prior expected breakpoint density is 11 per Morgan per offspring. The function `magicmask_impute` in the RABBIT, a combination of `magicmask` and `magicimpute`, was performed

~~~
magicmask_impute(sorghum_genofile,sorghum_design;
 isfounderinbred = true,
 model = “depmodel”,
 isphysmap = true,
 recomrate = 2.3,
 byfounder = 5,
 isrepeatimpute = true,
 tukeyfence = 2,
 iscorrectfounder = true,
 isspacemarker = true,
 skeletonsize = nothing,
)
~~~

where `model` is set to “depmodel” since the sorghum is a homozygous population; general default “jointmodel” was used for the rice F2 and the apple CP. The opitons `isphysmap` and `recomrate` were used to obtained initial genetic map before inferring inter-marker distances. The options `likeparam`, `threshlikeparam`, and `softhreshlikeparam` are set to the default values, where only `offspringerror` is relevant for “depmodel”. The option `byfounder` specifies the upper bound block size 5, since there are 10 founders being completely missing. The option `skeletonsize = nothing` indicates the number of skeleton markers is internally determined by the number of different genetic positions in each chromosome. A new `sorghum_genofile2` was output.

We then run function `magicreconstruct` in the RABBIT

~~~
 magicreconstruct(sorghum_genofile2,sorghum_design;
 isfounderinbred = true,
 hmmalg = “forwardbackward”,
)
~~~

where `hmmalg` denotes the forward backward algorithm for haplotype reconstruction. The resulting posterior ancestral haplotype probabilities were save in a text file `ancestryfile`.

~~~
Lastly, we performed QTL mapping using the statgenMPP package.
sorghum_mqm <- selQTLMPP(sorghum_mpp,
  trait = height, threshold = 5, maxCofactors = NULL,
    computeKin = true, parallel = TRUE)
~~~

where object `sorghum_mpp` was produced from `ancestryfile`, `trait` denotes the mean height data between 2013 and 2014, `maxCofactors` denotes the maximum number of co-factors to include in the model, `threshold` denotes the threshold of −*log*10 p-value for detecting a QTL, and `computeKin` denotes whether to calculate chromosome specific kinship matrices and include them in the model.

## Supplementary figures

**Figure S1.**
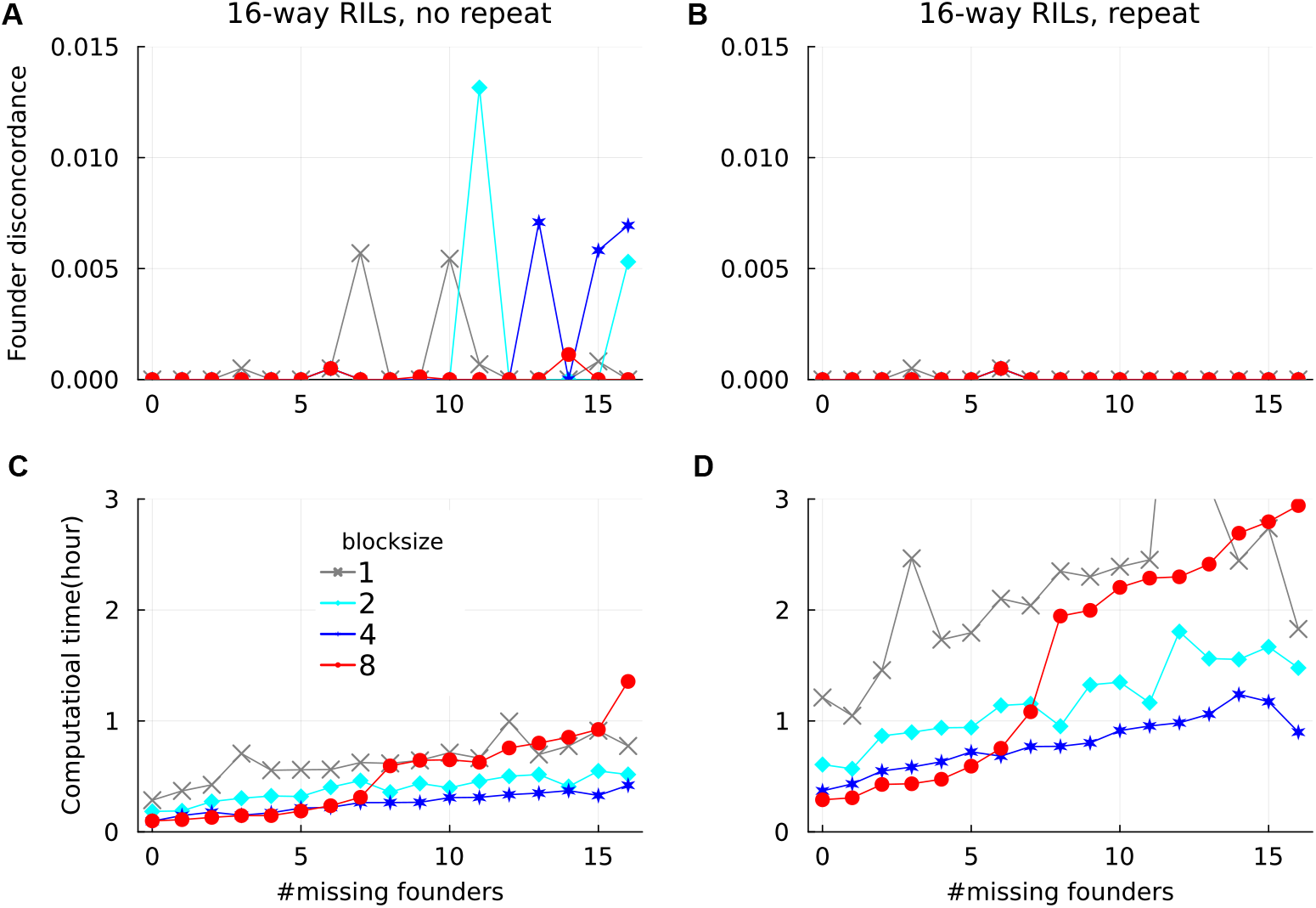
Founder imputation in the simulated 16-way RILs with population size 500. The x-axis denotes the number of founders with all genotypes being missing.

**Figure S2.**
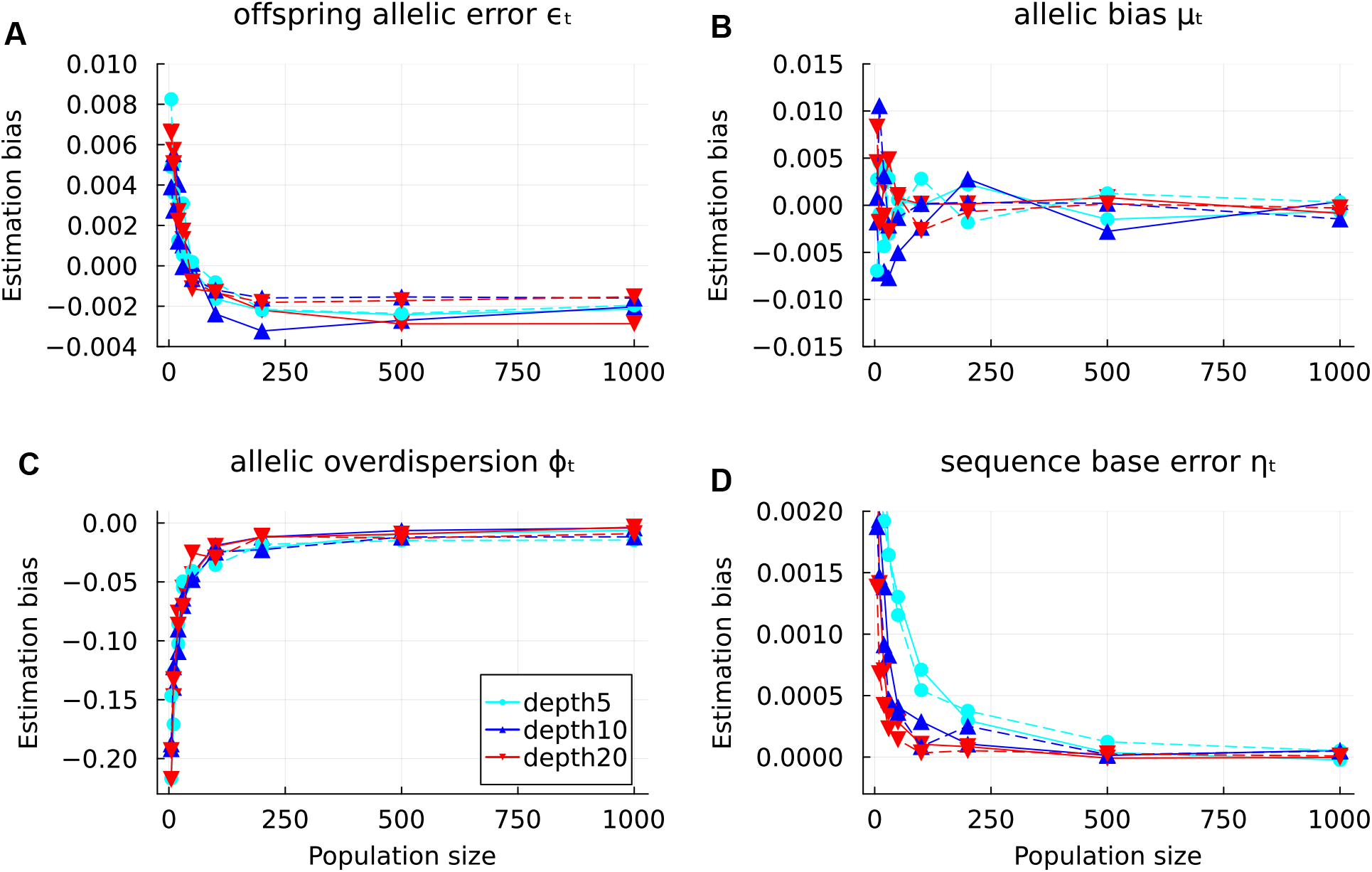
Estimation bias for the error rates in the simulated F2 (dashed lines) and CP (solid lines).

**Figure S3.**
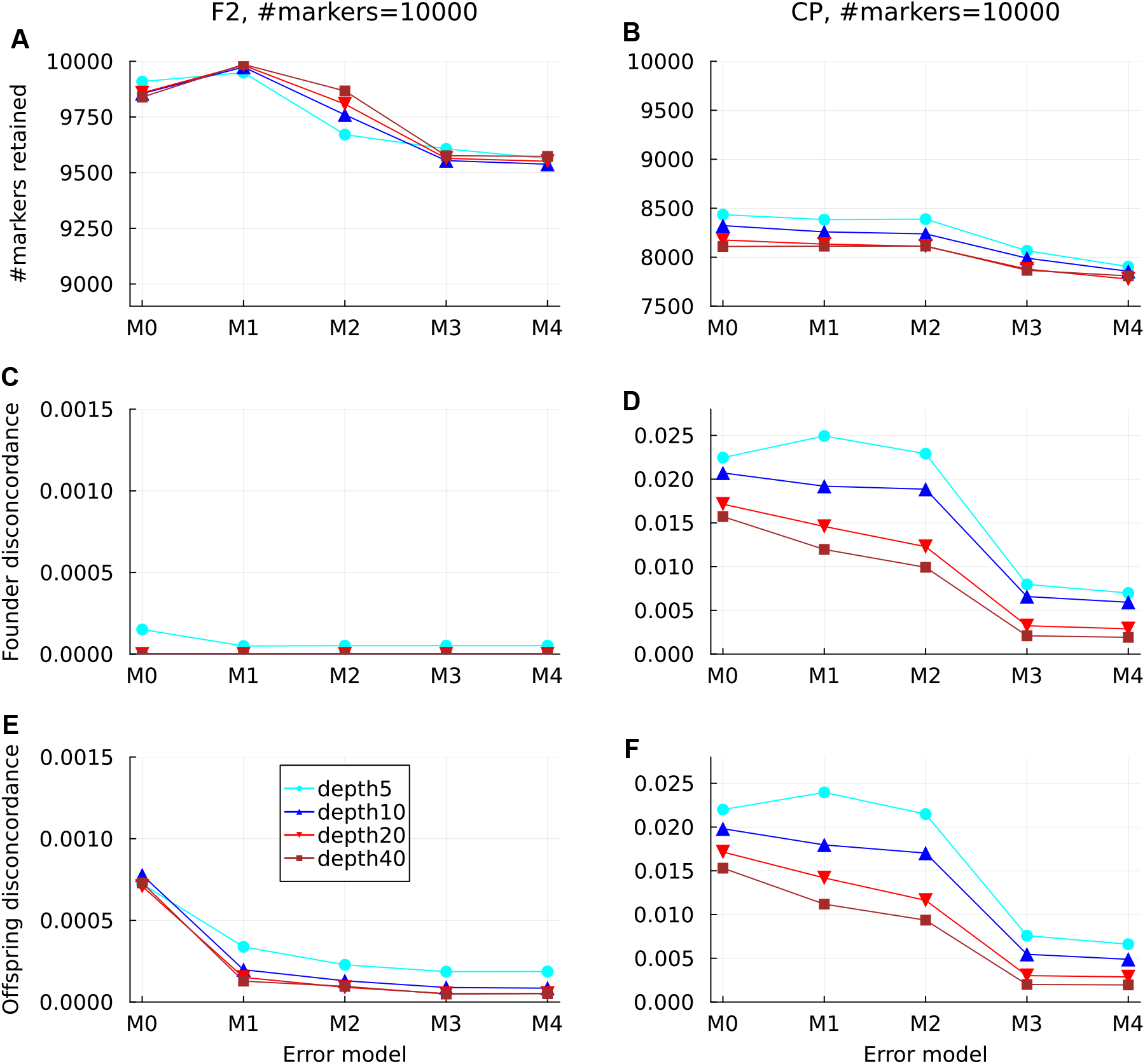
Genotype estimation in simulated F2 and CP without the founder error correction. The results were obtained with the default soft- and hard-threshold-based marker deletion.

**Figure S4.**
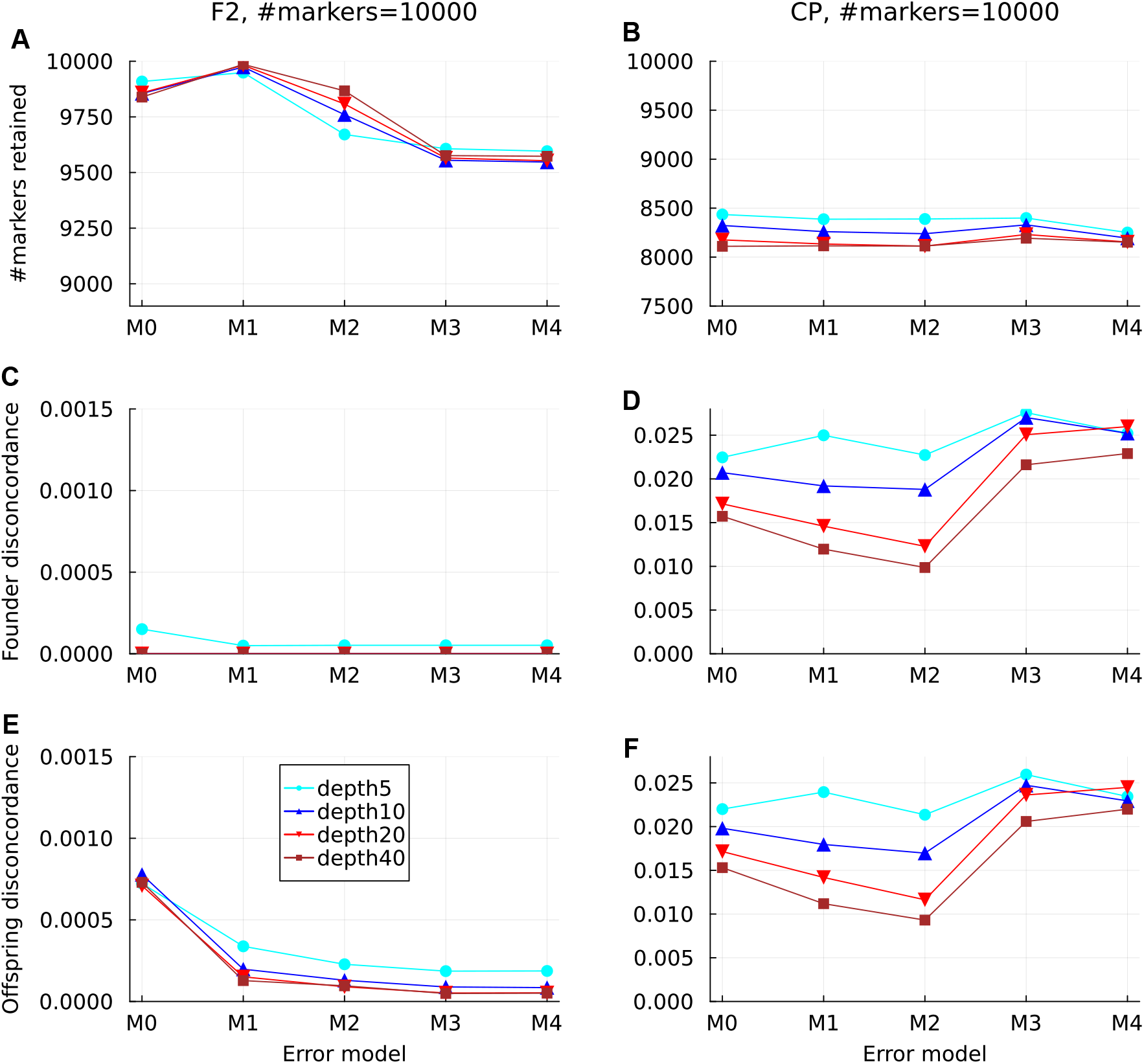
Genotype estimation in simulated F2 and CP with no founder error correction and no soft-threshold-based marker deletion. The results were obtained with the default hard-threshold-based deletion.

**Figure S5.**
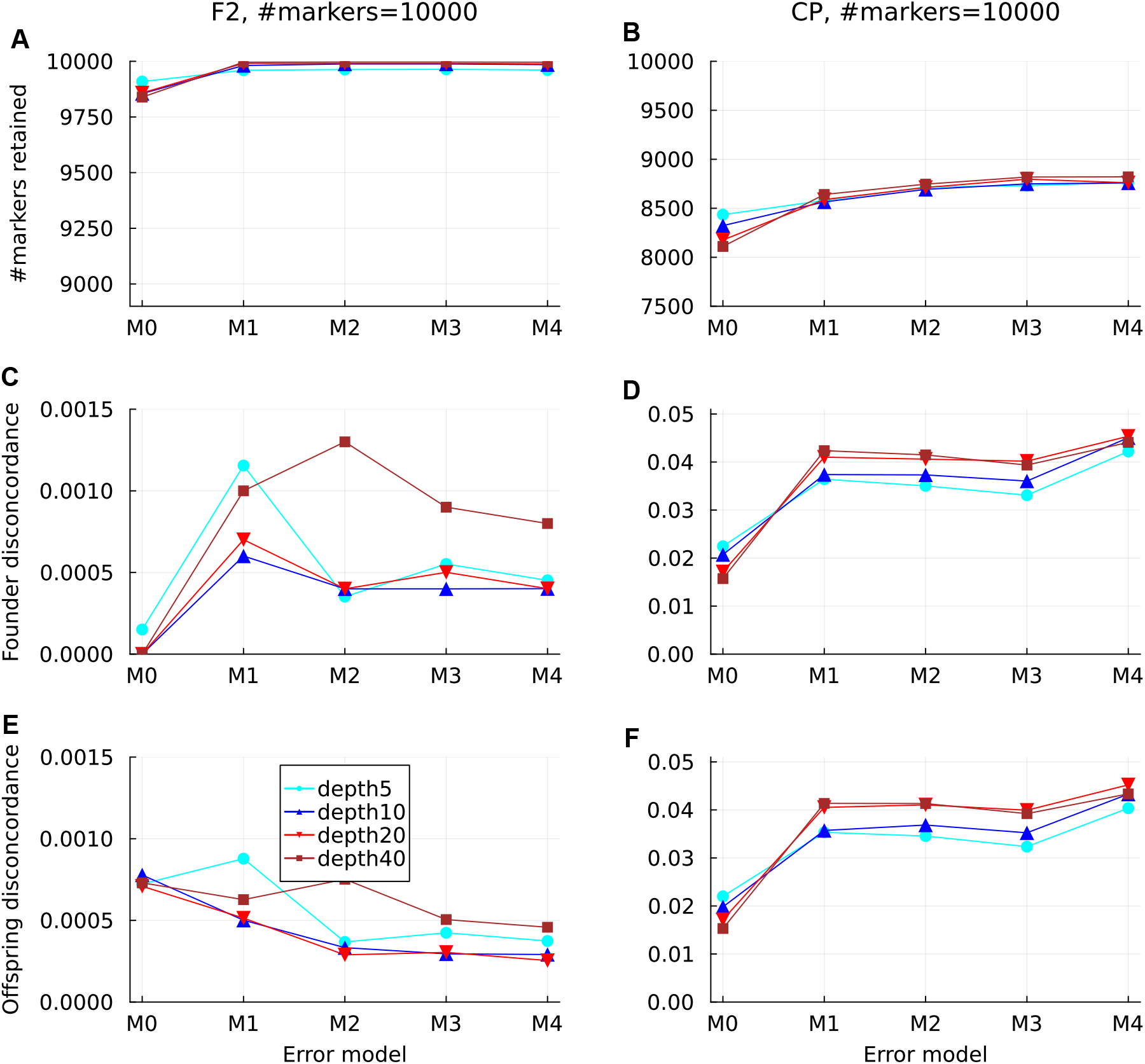
Genotype estimation in simulated F2 and CP with no founder error correction and no soft- or hard-threshold-based deletion.

**Figure S6.**
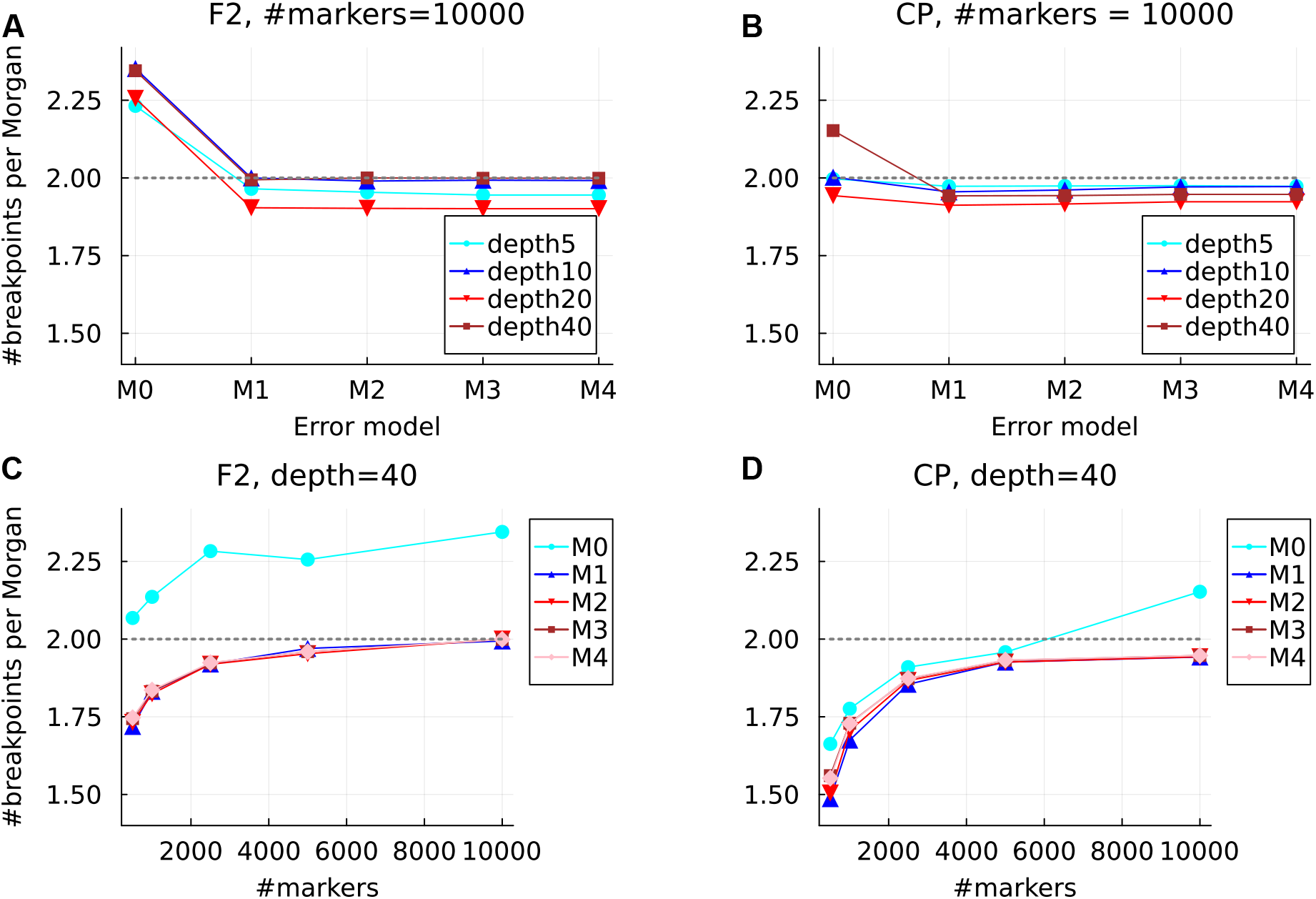
Estimated breakpoint density from the imputed genotypic data in the simulated F2 and CP. The error models are defined in Table 1. The horizontal dotted lines denote the prior number of breakpoints per Morgan per offspring.

**Figure S7.**
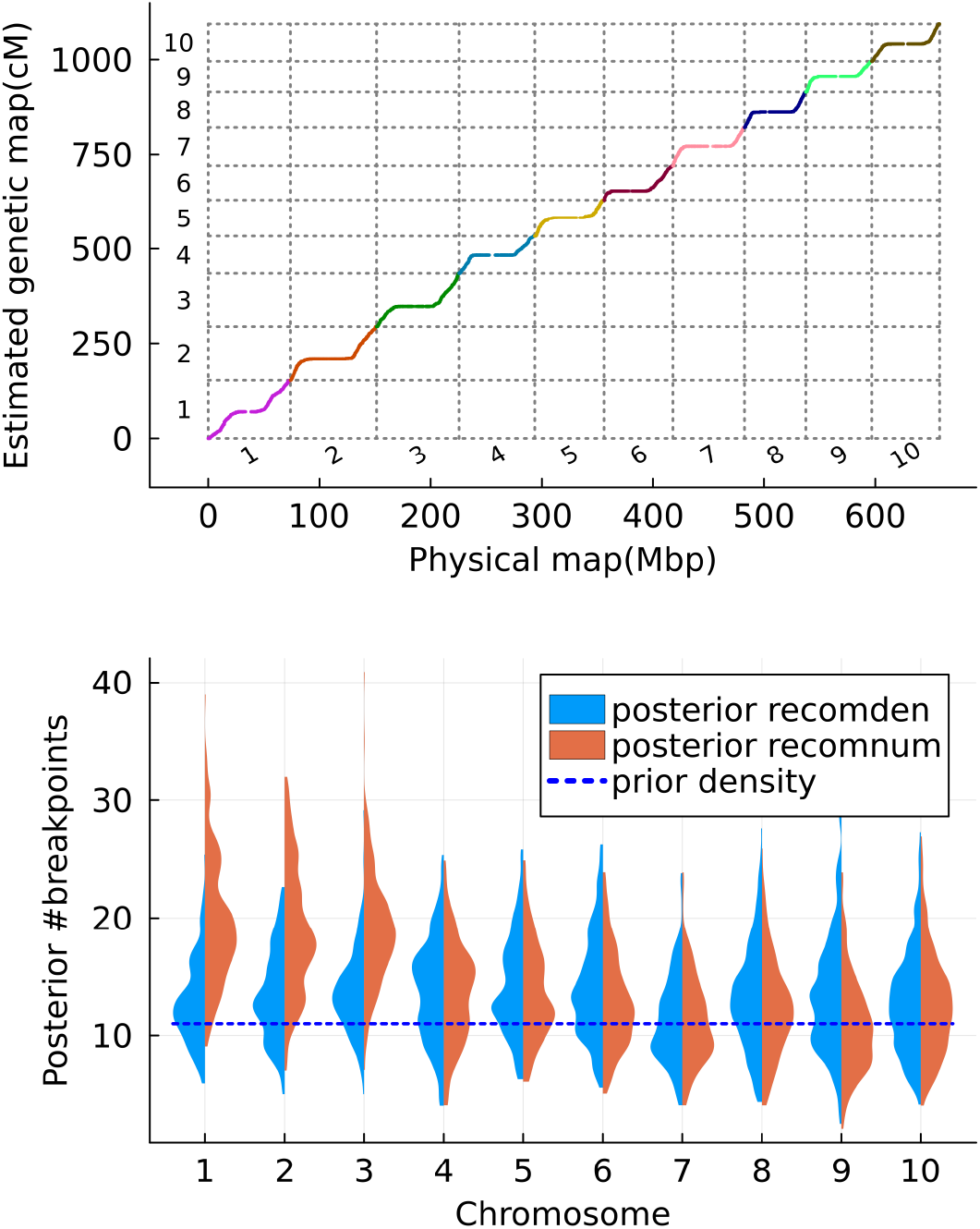
Estimated genetic map and number of breakpoints per chromosome in the real sorghum MAGIC. In the bottom panel, “recomnum” denotes the number of breakpoints per offspring, and “recomden” denotes the number of breakpoints per Morgan per offspring.

**Figure S8.**
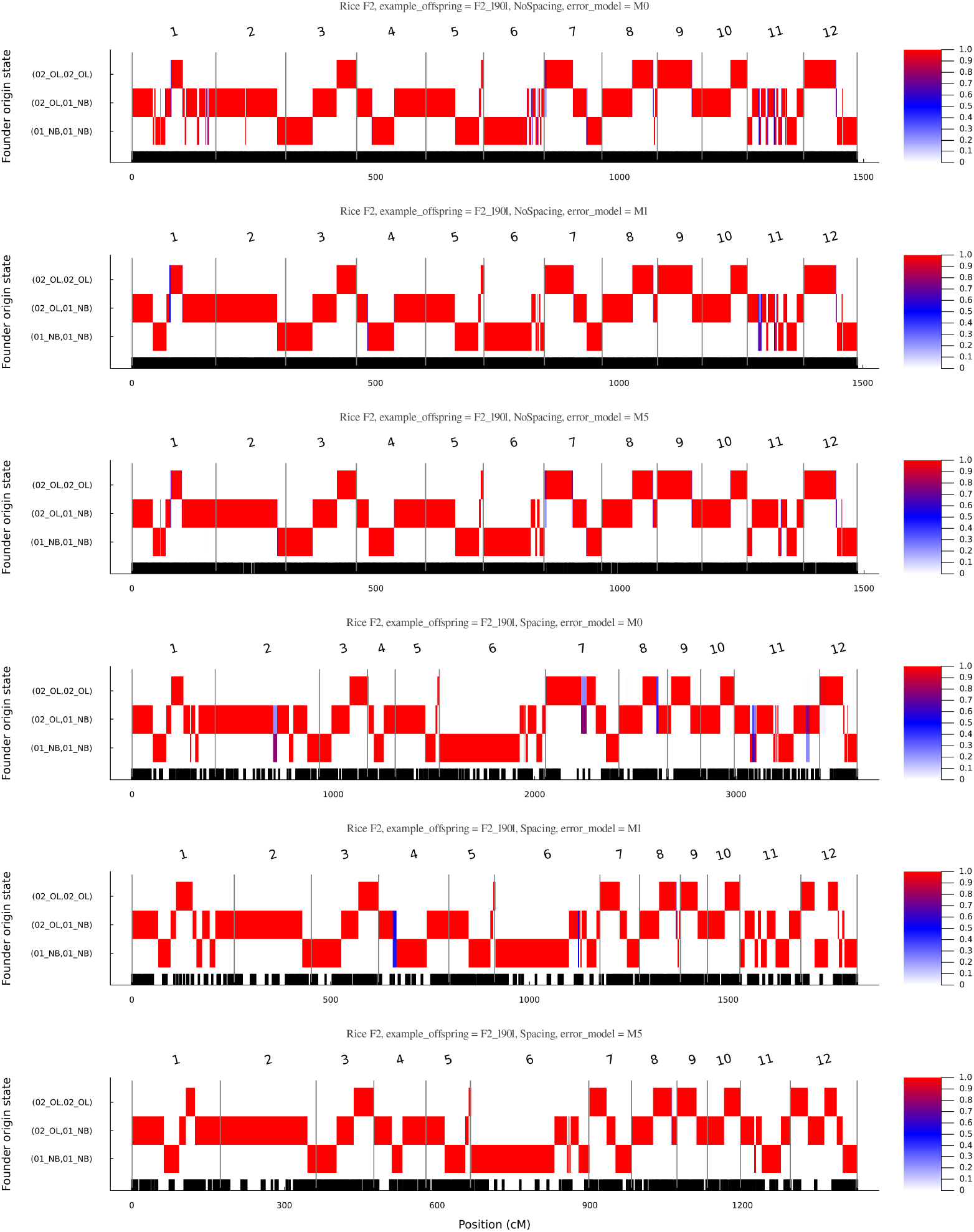
Posterior ancestral probabilities for an example offspring in the real rice F2. The first three panels refer to the results without inferring inter-marker distances, and the rest panels refer to the results with inferring distances.

**Figure S9.**
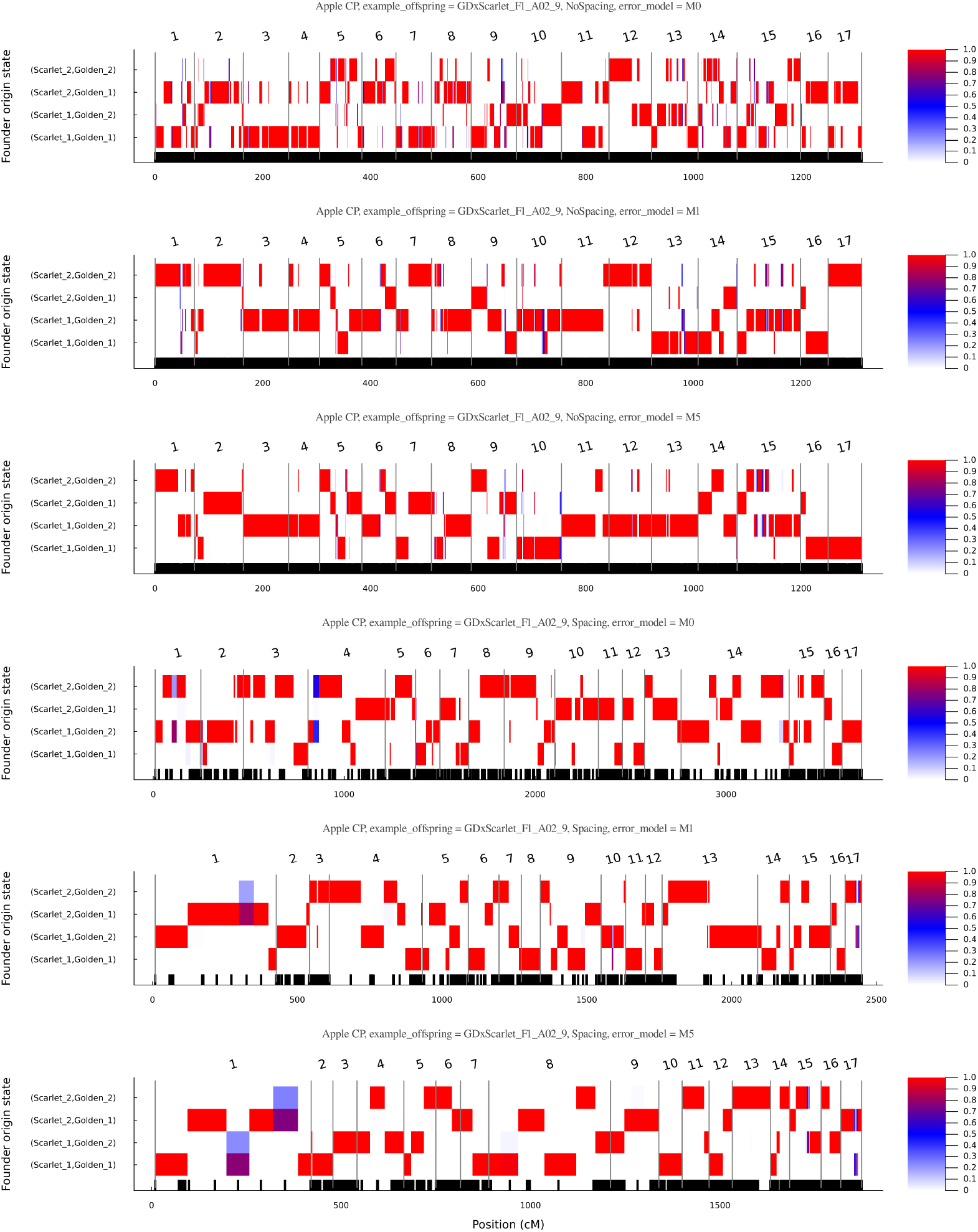
Posterior ancestral probabilities for an example offspring in the real apple CP. Note that the ordering of the two homologs for each parent in each chromosome is non-identifiable and thus the two homologs are arbitrarily labeled.

**Figure S10.**
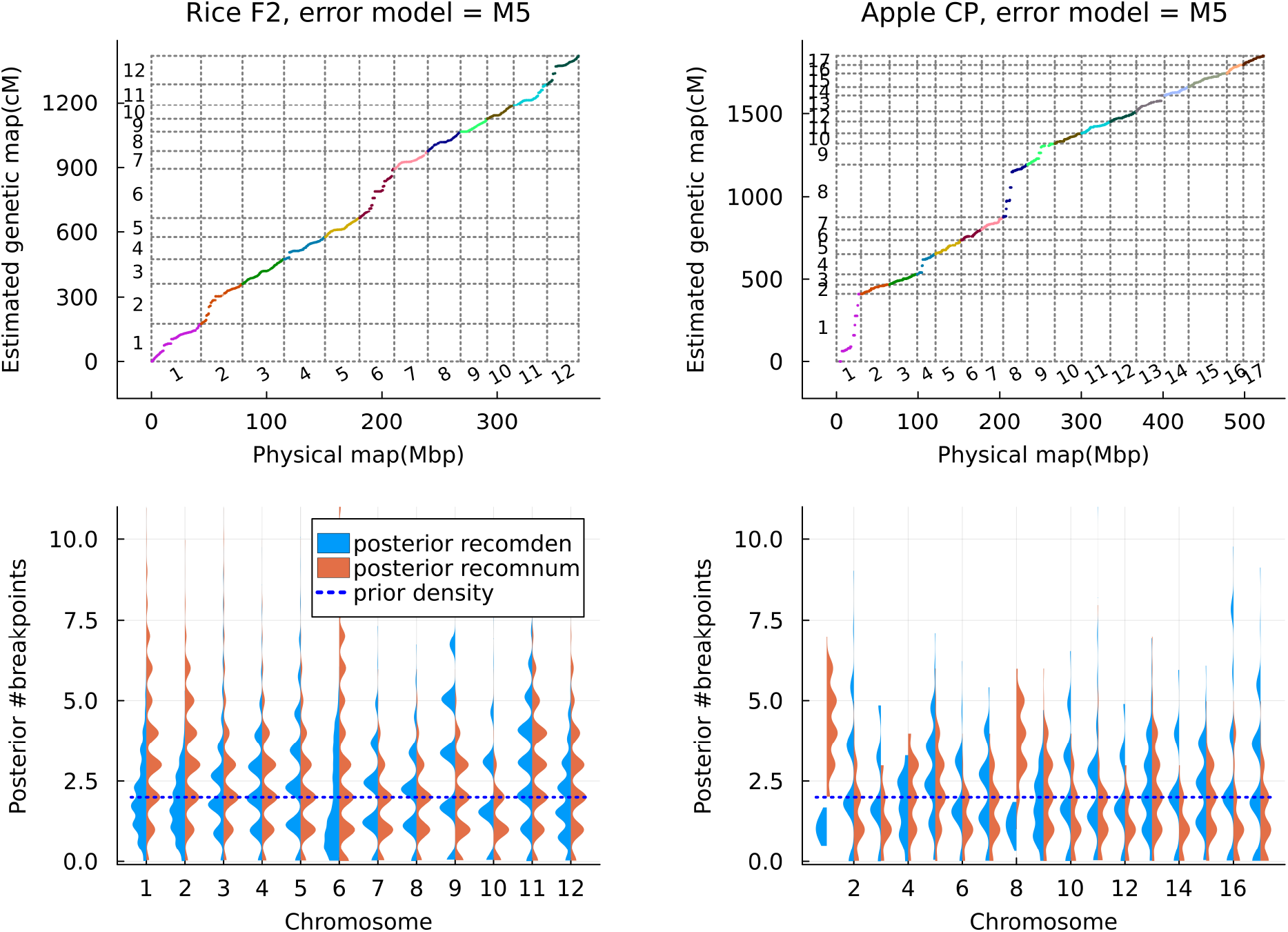
Estimated genetic map and number of breakpoints per chromosome in the real rice F2 and apple CP. In the bottom panels, “recomnum” denotes the number of breakpoints per offspring, and “recomden” denotes the number of breakpoints per Morgan per offspring.

**Figure S11.**
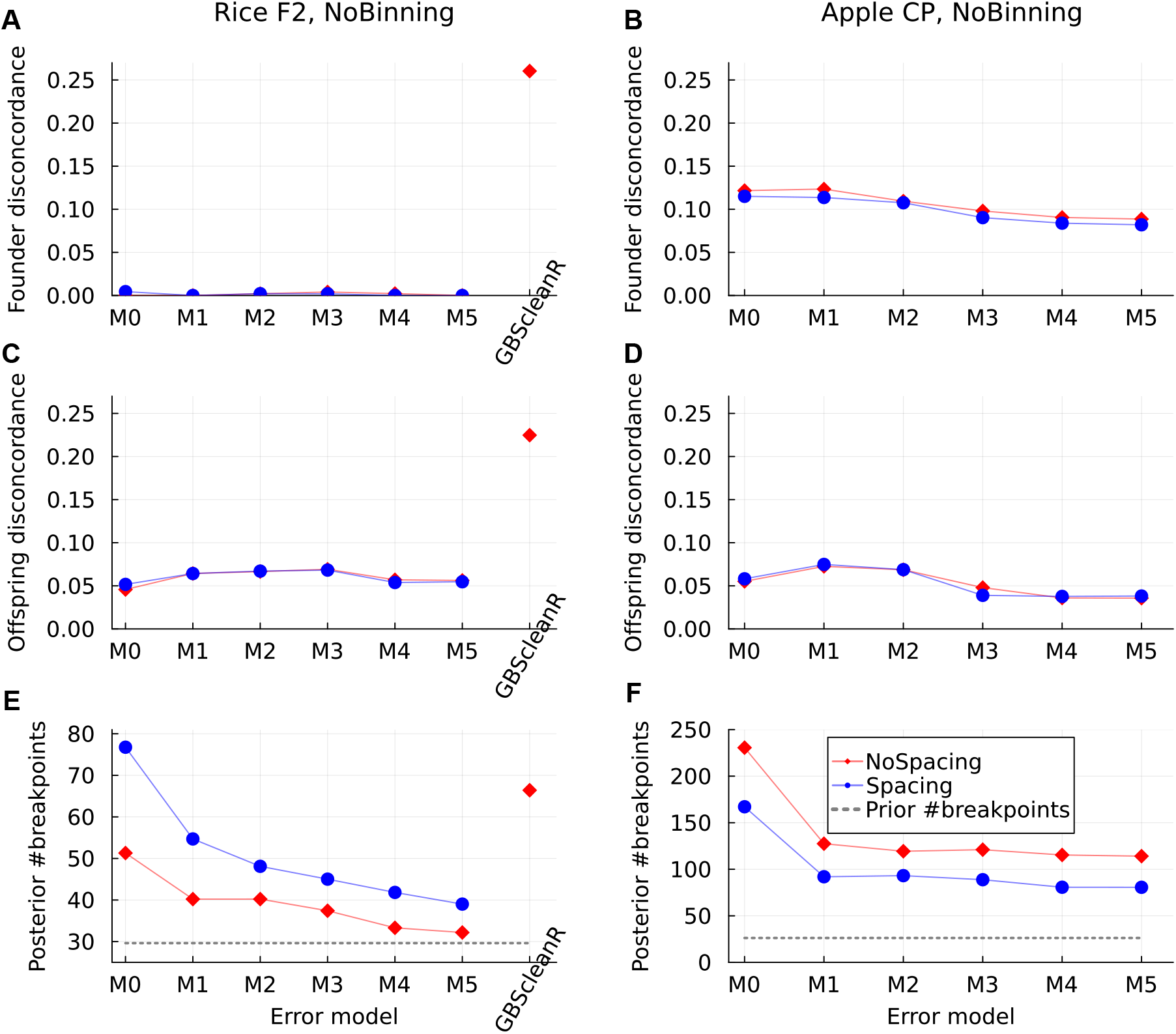
Founder imputation in the real F2 and CP without marker binning during imputation.

